# Deep Active Inference and Scene Construction

**DOI:** 10.1101/2020.04.14.041129

**Authors:** R. Conor Heins, M. Berk Mirza, Thomas Parr, Karl Friston, Igor Kagan, Arezoo Pooresmaeili

**Affiliations:** Department of Collective Behaviour, Max Planck Institute for Animal Behavior, D-78547 Konstanz, Germany; Perception and Cognition Group, European Neuroscience Institute;University Medical Centre Göttingen, Göttingen, Germany; Leibniz Science Campus “Primate Cognition”, Göttingen, Germany; Wellcome Centre for Human Neuroimaging, University College London, London, UK; Department of Neuroimaging, Institute of Psychiatry, Psychology and Neuroscience, King’s College London, London, UK; The NIHR Maudsley Biomedical Research Centre (BRC) at South London; Maudsley NHS Foundation Trust; the Institute of Psychiatry, Psychology and Neuroscience, King’s College London, London, UK; Decision and Awareness Group, German Primate Centre (DPZ), Göttingen, Germany

## Abstract

Adaptive agents must act in intrinsically uncertain environments with complex latent structure. Here, we elaborate a model of visual foraging – in a hierarchical context – wherein agents infer a higher-order visual pattern (a ‘scene’) by sequentially sampling ambiguous cues. Inspired by previous models of scene construction – that cast perception and action as consequences of approximate Bayesian inference – we use active inference to simulate decisions of agents categorizing a scene in a hierarchically-structured setting. Under active inference, agents develop probabilistic beliefs about their environment, while actively sampling it to maximise the evidence for their internal generative model. This approximate evidence maximization (i.e. self-evidencing) comprises drives to both maximise rewards and resolve uncertainty about hidden states. This is realised via minimization of a free energy functional of posterior beliefs about both the world as well as the actions used to sample or perturb it, corresponding to perception and action, respectively. We show that active inference, in the context of hierarchical scene construction, gives rise to many empirical evidence accumulation phenomena, such as noise-sensitive reaction times and epistemic saccades. We explain these behaviours in terms of the principled drives that constitute the *expected free energy*, the key quantity for evaluating policies under active inference. In addition, we report novel behaviours exhibited by these active inference agents that furnish new predictions for research on evidence accumulation and perceptual decision-making. We discuss the implications of this hierarchical active inference scheme for tasks that require planned sequences of information-gathering actions to infer compositional latent structure (such as visual scene construction and sentence comprehension). Finally, we propose experiments to contextualise active inference in relation to other formulations of evidence accumulation (e.g. drift-diffusion models) in tasks that require planning in uncertain environments with higher-order structure.

## 1 Introduction

Our daily life is full of complex sensory scenarios that can be described as examples of ‘scene construction’ (Hassabis & Maguire, 2007; Zeidman, Lutti, & Maguire, 2015; Mirza, Adams, Mathys, & Friston, 2016). In its most abstract sense, scene construction describes the act of inferring a latent variable (or ‘scene’) given a set of (potentially ambiguous) sensory cues. Sentence comprehension is a prime example of scene construction: individual words are inspected in isolation, but after reading a sequence one is able to abduce the overall meaning of the sentence that the words are embedded within (Tanenhaus, Spivey-Knowlton, Eberhard, & Sedivy, 1995; Narayanan, Jurafsky, et al., 1998). This can be cast as a form of hierarchical inference in which low-level evidence (e.g. words) is actively accumulated over time to support disambiguation of high-level hypotheses (e.g. possible sentence meanings). This dialogue between different layers of inference (e.g. word identity ↔ sentence meaning) exemplifies the hierarchical nature of many inference scenarios encountered by agents navigating a complex world. In the current work, we explore this hierarchical dialogue while parametrically varying the uncertainty present at low- and high-levels of inference. The analogue of this - in the reading example - would be the lexico-semantic processing incurred by understanding a messily-handwritten sentence: as words are successively read, comprehension of the sentence meaning gets easier. This updated, high-level belief may then ‘percolate down’ to support word identification, resulting in resolution of uncertainty at the individual word-level (Dilkina, McClelland, & Plaut, 2010; Evans, Ralph, & Woollams, 2012).

We investigate this sort of hierarchical belief-updating by modelling scene construction in a visual foraging task, where visual stimuli are actively sampled with saccadic eye movements in order to accumulate information and categorise accurately (Yarbus, 1967; Jóhannesson, Thornton, Smith, Chetverikov, & Kristjánsson, 2016; Mirza et al., 2016; Yang, Lengyel, & Wolpert, 2016; Ólafsdóttir, Gestsdóttir, & Kristjánsson, 2019). This is an example of active vision, an area of research focusing on the closed causal loop that links visual processing to the actions that result in visual sensation. For example, animals like primates can actively alter their visual sensorium by moving their eyes to sample new visual data (Rolfs, 2015; Itti & Baldi, 2009; Parr & Friston, 2017a; Yang, Wolpert, & Lengyel, 2018; Gottlieb & Oudeyer, 2018). As mentioned in the earlier example, the momentary contents of visual fixation are often contaminated with noise. This noise can have either internal sources (e.g. neural noise) or be an intrinsic property of visual stimuli themselves. In the context of scene construction, such sensory uncertainty can limit the ability of individual cues to support inference about the overarching visual scene. These limitations can nevertheless be partially overcome if the agent has prior knowledge, which can be built into the agent’s belief structure, innately or as a function of previous experience. While there is an enormous body of literature on the resolution of uncertainty with prior information in simple scenarios (Trueswell, Tanenhaus, & Garnsey, 1994; Körding & Wolpert, 2004; Stocker & Simoncelli, 2006; Girshick, Landy, & Simoncelli, 2011), relatively little research has examined interactions between sensory uncertainty and prior information in the context of a dynamic, active vision task like visual foraging or scene construction (with notable exceptions: e.g. (Quétard et al., 2016)).

In this work we cast visual scene construction as an inference problem, in which action and perception both serve to maximise evidence for an internal generative model of sensory data and its hidden causes. Building on a previous Bayesian formulation of scene construction, we use *active inference* to model perception and active evidence accumulation in a hierarchical scene construction task (Friston, Adams, Perrinet, & Breakspear, 2012; Friston, Samothrakis, & Montague, 2012; Mirza et al., 2016; Friston, FitzGerald, Rigoli, Schwartenbeck, & Pezzulo, 2017).

Using a generic model of a hierarchical visual foraging scenario, we explore the relationship between sensory uncertainty and prior information and how they jointly determine behaviour and belief dynamics under active inference. The (sometimes counterintuitive) results of our simulations invite new perspectives on active sensing and hierarchical inference, which we discuss in the context of experimental design for both visual foraging experiments and perceptual decision-making tasks more generally. We also examine the model’s behaviour in terms of the tension between instrumental (or utility-driven) and exploratory (epistemically-driven) drives, and how active inference explains both apparently-distinct kinds of value by evaluating policies with respect to a single pseudo-’value function’: an expected bound on surprise called the *expected free energy*.

The rest of this paper is organised as follows: first, we will motivate the task by introducing active vision and scene construction, relating the problem faced in these scenarios to the widely-studied phenomenon of ‘evidence accumulation’ in cognitive neuroscience. This discussion of evidence accumulation then leads us to introduce a class of psychophysical stimuli known as ‘random dot motion’ stimuli (RDMs). We will use RDMs as sensory stimuli in our formulation of hierarchical scene construction. We then summarise the theoretical background of active inference and the free energy principle, highlighting the utility of active inference as a generic model of adaptive behaviour in complex systems. Next, we detail the Markov Decision Process we use to simulate perception and action in active inference agents and the dynamics of the environment. Having specified a Markovian generative model for the scene construction task, we then report the differential effects of sensory uncertainty and prior belief strength on several aspects of evidence accumulation in active inference simulations. These computational demonstrations motivate our conclusion, where we discuss the implications of this work for experimental studies of active sensing and evidence accumulation under uncertainty.

## 2 Active Vision and Inference

Active vision research emphasises the fact that in many artificial and biological visual systems, sensing apparati are actively deployed in order to gather visual information. More generally, active sensing approaches study the embodied, self-initiated aspect of perception in different sensory modalities (Bajcsy, 1988; König & Luksch, 1998). This is consistent with the so-called ‘4Es’ (Embodied, Embedded, Enacted, Extended) framework for studying cognition (Rowlands, 2010). This is a concept in the cognitive and mind sciences that has emerged over the last few decades, and emphasises the deep connection between cognitive operations (perception, decision-making) and the physical realities (e.g. morphology of the sense organs) that constrain biological agents.

Under many active sensing accounts, agents are thought to entertain (explicitly or not) hypotheses about hidden states of the world that give rise to their sense data (Buckley, Kim, McGregor, & Seth, 2017). These world-states are external and thus unknown to the agent, but the agent can nevertheless predict their sensory consequences given a sufficient understanding of how world-states evince observations. In a Bayesian sense, the agent’s beliefs about the world and how it causes sensations comprise what’s known as a *generative model*. This leads to the idea of perception as problem of statistical inference, wherein sensory data is inverted by means of the agent’s generative model to estimate the most likely causes of sensations in the world. This conception originates in the ideas of Helmholtz and was developed by others in the machine learning and cognitive science communities (von Helmholtz, 1867; Gregory, 1980; Hinton & Zemel, 1994; Dayan, Hinton, Neal, & Zemel, 1995; Knill & Pouget, 2004; Clark, 2013). Given this formulation of perception as inference, active sensing is then be understood to be a form of hypothesis testing (e.g. ‘am I looking at a dog or a cat?’ or ‘where’s Waldo?’), wherein evidence supporting various competing explanations for sensory data is actively gathered through the deployment of sense organs. In the case of vision, for example, uncertainty about the environmental causes of visual stimulation motivates inspection of the visual world with eye movements (Itti & Baldi, 2009; Friston, Adams, et al., 2012; Yang et al., 2016; Mirza et al., 2016; Parr & Friston, 2017a, 2017b).

The particular neuronal implementations underlying active visual processes and its consequences for perception and action have been the subject of a wealth of cognitive neuroscience research (Foley, Jangraw, Peck, & Gottlieb, 2014; Adams, Aponte, Marshall, & Friston, 2015; Baranes, Oudeyer, & Gottlieb, 2015; Daddaoua, Lopes, & Gottlieb, 2016; Yang et al., 2016; Foley, Kelly, Mhatre, Lopes, & Gottlieb, 2017; Mirza, Adams, Mathys, & Friston, 2018; Parr, Mirza, Cagnan, & Friston, 2019). It is also worth noting the conceptual affinity this research has to investigations of related phenomena in neuroscience like attention, curiosity, and intrinsic motivations (Gottlieb, Lopes, & Oudeyer, 2016; Gottlieb & Oudeyer, 2018). As we will see later, active inference offers a Bayesian account of active sensing that simultaneously unites the related concepts of curiosity, epistemic motivation, and attention under the theoretical umbrella of inference (Friston et al., 2015; Friston, FitzGerald, et al., 2017; Friston, Lin, et al., 2017; Schwartenbeck et al., 2019).

## 3 Perceptual Decision-Making and Evidence Accumulation

Seminal work on non-human primates in the 1980s and 1990s laid the groundwork for a branch of neuroscience commonly referred to as ‘perceptual decision-making’ (Gold & Shadlen, 2007). Subsequent theory attempted to model the process underlying perceptual decisions as the dynamics of a decision variable whose instantaneous value is related to the amount of evidence currently available to discriminate a stimulus’ identity - hence the term *evidence accumulation* (Ratcliff & Rouder, 1998; Bogacz, Brown, Moehlis, Holmes, & Cohen, 2006; Ratcliff & McKoon, 2008b). Properties of the decision variable’s dynamics have been related to experimental parameters like stimulus strength and the prior beliefs of the decision-maker about stimulus preponderance, among others (Bitzer, Park, Blankenburg, & Kiebel, 2014; Fard, Park, Warkentin, Kiebel, & Bitzer, 2017). Upon reaching an ‘evidence-bound’ or decision threshold, the decision variable then triggers the perceptual recognition or decision, which is often measured as the choice of one out of a discrete set of response options, with each option reporting recognition of a particular stimulus.

Of relevance to the current investigation is a string of studies of evidence accumulation in the context of visual motion processing (Newsome & Paré, 1988; Salzman, Britten, & Newsome, 1990; Newsome, Britten, & Movshon, 1989; Britten, Shadlen, Newsome, & Movshon, 1992, 1993; Shadlen & Newsome, 1994, 1996, 2001). The prototypical stimulus used in these experiments is the ‘random dot motion’ display or RDM (Nakayama & Silverman, 1984). An RDM pattern consists of a small patch of dots whose correlated displacement over time gives rise to the perception of apparent directed motion (see Figure 1). By manipulating the proportion of dots moving in the same direction, the apparent direction of motion can be made more or less difficult to discriminate. This discriminability is usually operationalised as a single *coherence* parameter, which defines the percentage of dots that appear to move in a common direction. The remaining non-signal (or ‘incoherent’) dots are usually designed to move in random independent directions. This coherence parameter thus becomes a simple proxy for sensory uncertainty in motion perception: manipulating the coherence of RDM patterns has well-documented effects on behavioural measures of performance such as reaction time and discrimination accuracy, with increasing coherence leading to faster reaction times and higher accuracy (Palmer, Huk, & Shadlen, 2005). In the following section we synthesise the current and previous sections to present our generic model of hierarchical scene construction, which requires the active accumulation of individual sensory stimuli (here, RDM patterns) in the service of scene construction.

**Figure 1:**
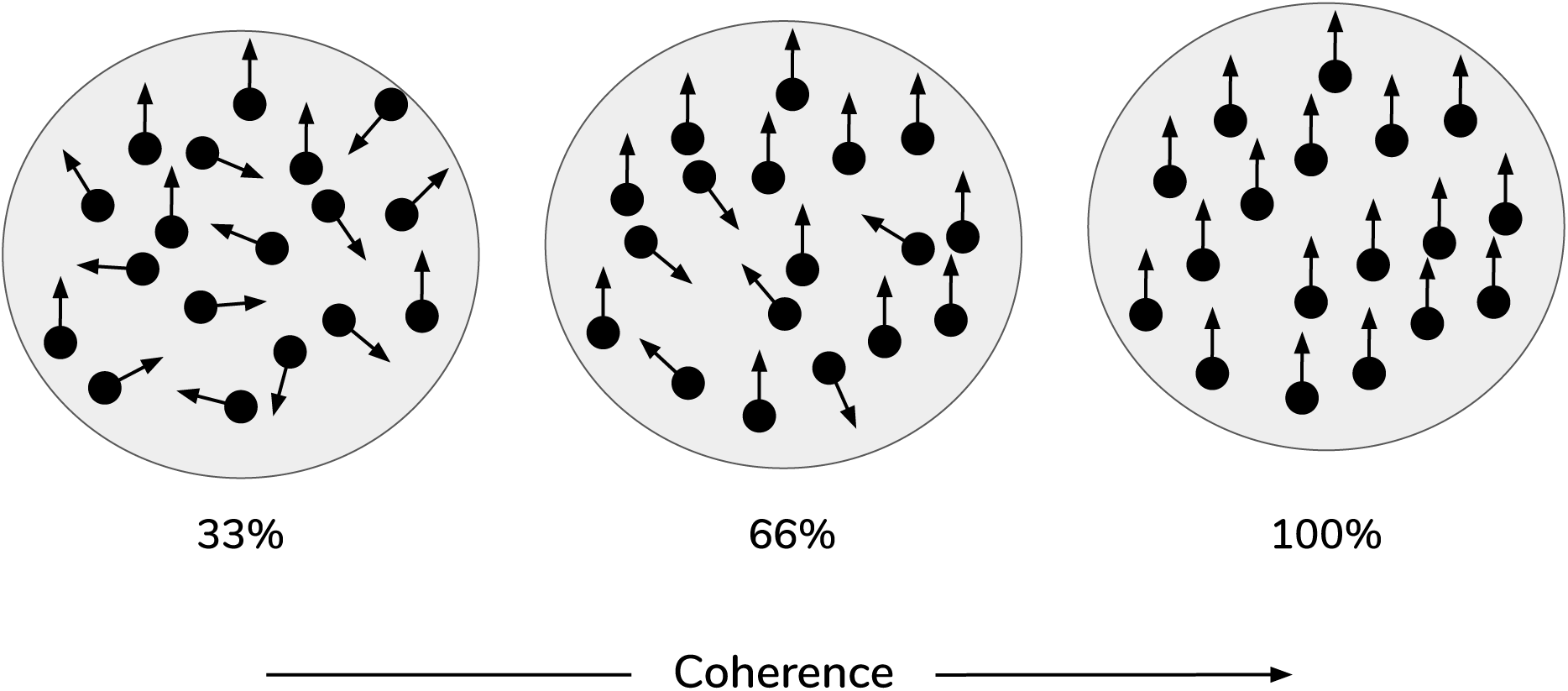
Random Dot Motion Stimuli (RDMs). Schematic of random dot motion stimuli, with increasing coherence levels (i.e. % percentage of dots moving upwards) from left to right.

## 4 Scene Construction with Random Dot Motion

We now describe an abstract scene construction task that will serve as the experimental context within which to frame our hierarchical account of active evidence accumulation. Inspired by a previous active inference model of scene construction introduced by (Mirza et al., 2016), here we invoke scene construction in service of a categorization game. In each trial of the task, the agent must make a discrete choice to report its belief about the identity of the ‘hidden scene.’ In the formulation by Mirza and colleagues, the scenes are represented by 3 abstract semantic labels: ‘flee,’ ‘feed,’ and ‘wait’ (see Figure 2). Each scene manifests as a particular spatial coincidence of two pictures, where each picture is found within a single quadrant in a 2 x 2 visual array. For example, the ‘flee’ scene is defined as when a picture of a cat and a picture of a bird occupy two quadrants lying horizontally adjacent to one other. The scene identities are also invariant to two spatial transformations: vertical and horizontal inversions. For example, in the ‘flee’ scene, the bird and cat pictures can be found in either in the top or bottom row of the 2 x 2 array, and they can swap positions; in any of these cases the scene is still ‘flee.’ The task requires active visual interrogation of the environment because quadrants must be *gaze-contingently* unveiled. That means, by default all quadrants are covered and their contents not visible; the agent must directly look at a quadrant in order to see its contents. This task structure and the ambiguous nature of the picture → scene mapping means that agents need to actively forage for information in the visual array in order to be certain about the scene in play.

**Figure 2:**
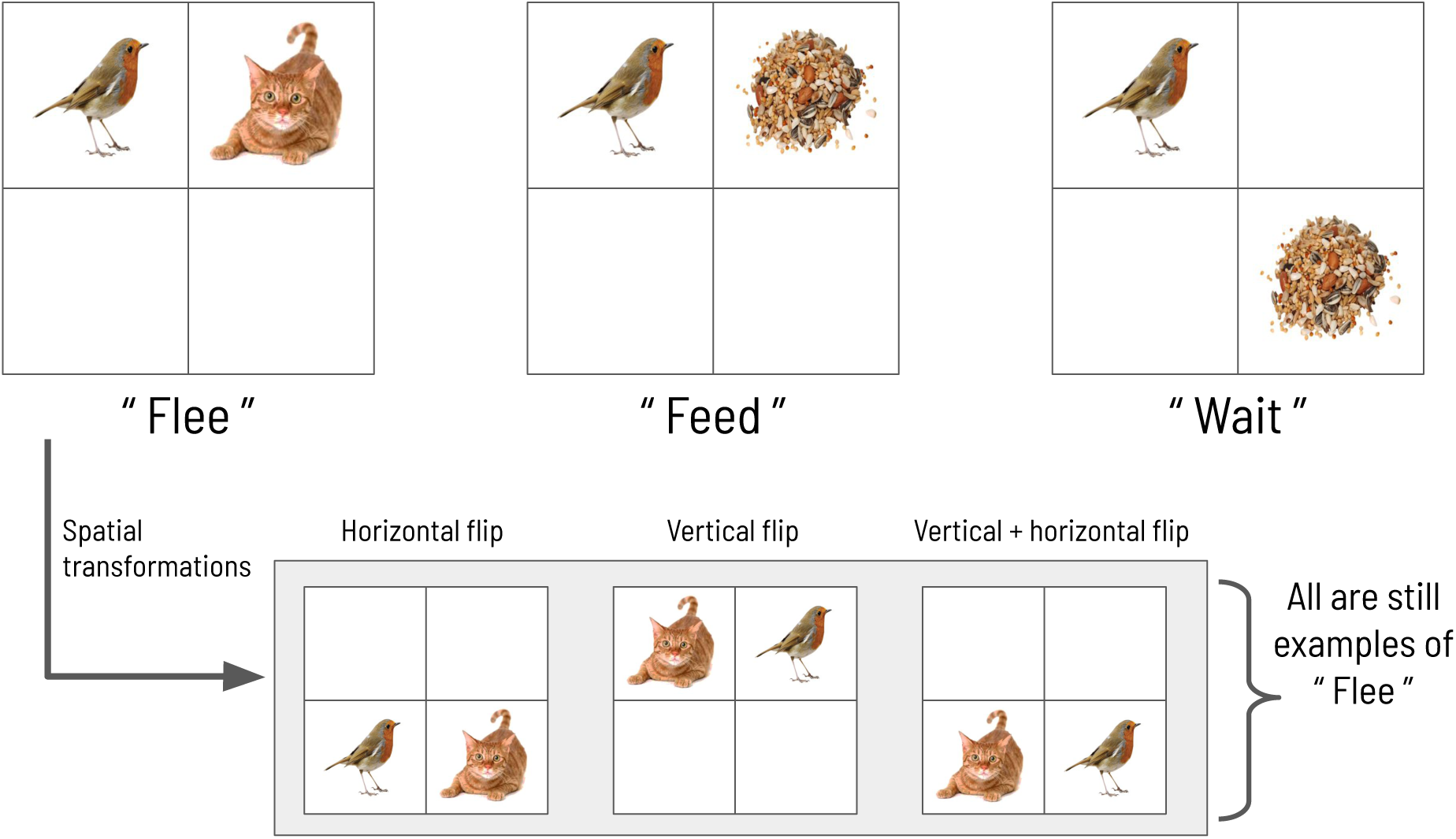
The scene configurations of the original formulation. The three scenes characterizing each trial in the original scene construction study, adapted with permission from (Mirza et al., 2016).

In the current work, scene construction is also treated as a categorization task, requiring the gaze-contingent disclosure of quadrants whose contents provide evidence for or against a particular scene. However, in the new task, the visual stimuli occupying the quadrants are RDMs that are distinguished by their distinct motion directions. Each RDM pattern is characterised by a primary direction of motion that is always one of the four cardinal directions: **UP, RIGHT, DOWN**, or **LEFT**. For example, in a given trial one quadrant may contain a motion pattern moving upwards, while another quadrant contains a motion pattern moving leftwards. These RDM stimuli are suitable for the current task because we can use the coherence parameter to tune motion ambiguity and hence sensory uncertainty. Applying this metaphor to the original task (Mirza et al., 2016): imagine that the bird and cat pictures are somehow made blurry, such that it becomes difficult to tell whether a given image is of a bird or a cat - this low-level uncertainty about individual images may then ‘carry forward’ to affect scene inference. An equivalent analogy might be found in the problem of inferring the meaning of a written sentence (the semantic equivalent of a ‘scene’) when the individual words are poorly written or illegible. In our case, the motion coherence of RDMs controls how easily an RDM of one direction can be confused with another direction - namely, a more incoherent dot pattern is more likely to be mistaken as a dot pattern moving in a different direction.

We also design the visual stimulus → scene mapping such that scenes are degenerate with respect to individual visual stimuli, as in the previous task (see Figure 3). There are four scenes, each one defined as the co-occurrence of two RDMs in two (and only two) quadrants of the visual array. The two RDMs defining a given scene move in perpendicular directions; the scenes are hence named: **UP-RIGHT, RIGHT-DOWN, DOWN-LEFT**, and **LEFT-UP**. Similar to the original scene construction formulation, discerning the direction of one RDM is not sufficient to disambiguate the scene; the player must observe two RDMs and discern their respective directions before being able to unambiguously infer scene identity. The task requires two nested inferences - one about the contents of the currently-fixated quadrant (e.g. ‘Am I looking at an **UP**-wards moving RDM’) and another about the identity about the overarching scene (e.g. is the scene ‘**UP-RIGHT**’?). During each trial, the agent can report its guess about the scene identity by choosing one of the four symbols that signify the scenes (see Figure 3) and doing so ends the trial. This concludes our narrative description of the experimental setup.

**Figure 3:**
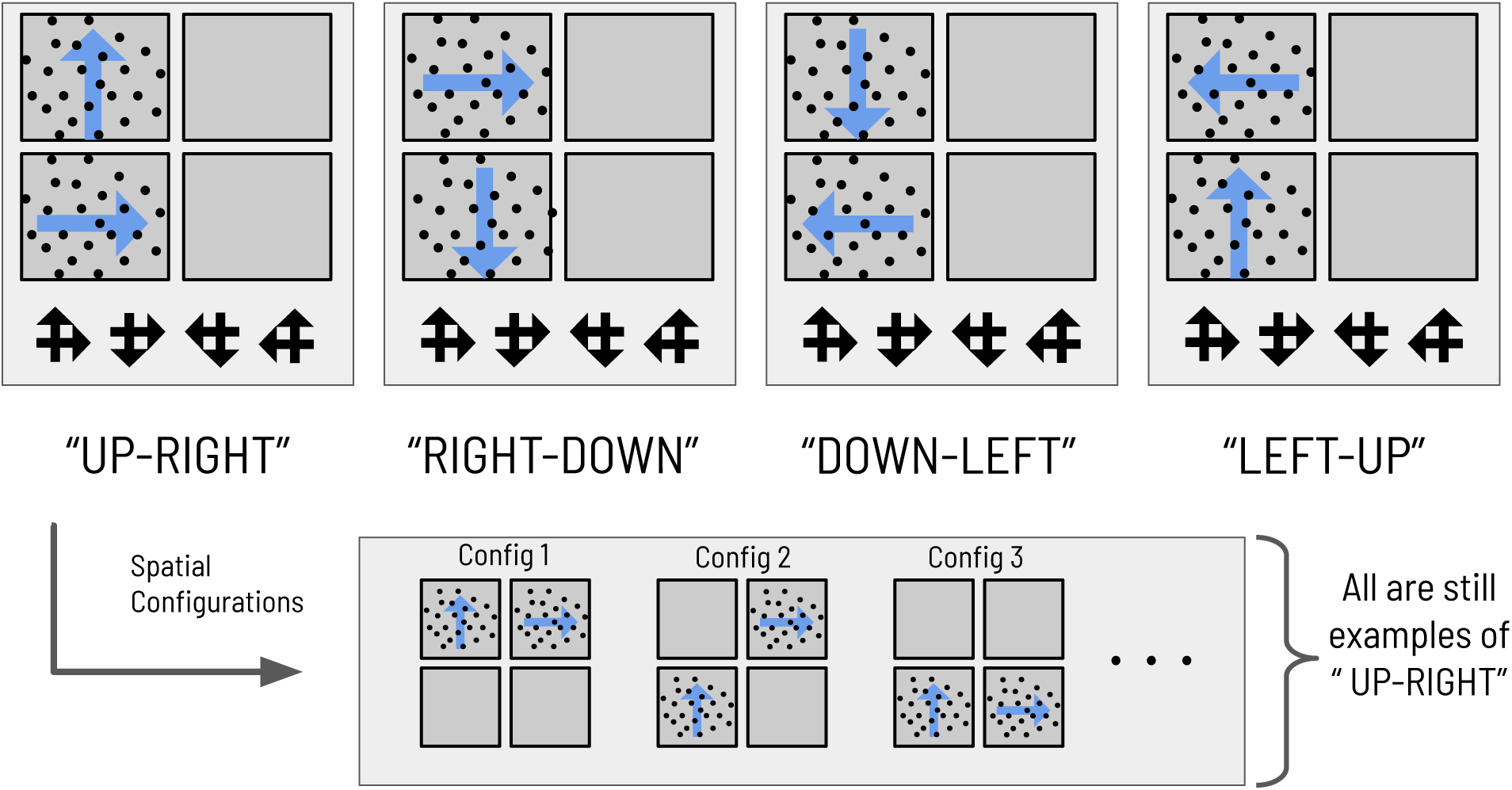
The mapping between scenes and RDMs. The mapping between the four abstract scene categories and their respective dot motion pattern manifestations in the context of the hierarchical scene construction task. As an example of the spatial invariance of each scene, the bottom right panels show two possible (out of 12 total) RDM configurations for the scene ‘**RIGHT-DOWN**’, where the two constitutive RDMs of that scene are found in exactly two of the four quadrants.

In the next section, we provide background on the Free Energy Principle and the ensuing active inference theory of Bayesian action and perception. With this theoretical background in mind, we will be in a position to formulate the scene construction task as a hierarchical Markov Decision Process that can be solved with active inference.

## 5 Free Energy Minimization and Active Inference

### 5.1 The thermodynamics of minimizing surprise

As a complex system persists over time, by definition it resists the external fluctuations that threaten its integrity; i.e., those that threaten dissolution, dissipation or decay. Said another way, the system can be described as forming probabilistic beliefs about the sources of variation in its world that are relevant to its continued existence. This equivalence, between a system’s failure to perish and the apparent drive to persist and model the milieu, is the philosophical underpinning of the Free Energy Principle (or FEP) (Friston, 2019). It is often expressed in terms of the information theoretic notion of surprise and its approximate minimization in Bayesian inference schemes (Beal, 2004; Friston, 2012; Ramstead, Badcock, & Friston, 2018; Kirchhoff, Parr, Palacios, Friston, & Kiverstein, 2018). Originally conceived as a unifying explanation for the structure and function of the cerebral cortex (Friston, 2005; Friston & Kiebel, 2009; Friston, 2009, 2010), FEP has matured into a framework used to describe the fundamental drive of complex, adaptive systems to maintain their internal states within bounds that are conducive to survival.

The concept underpinning the so-named principle is the *variational free energy*, a function(al) that adaptive systems must minimise as a consequence of maintaining statistical integrity in the face of external fluctuations. An intuitive grip on the underlying Bayesian mechanics emerges when describing variational free energy as an upper bound on *surprise*, where surprise is the log probability of sensory perturbations *y*, given some model *m* of how these perturbations are generated:

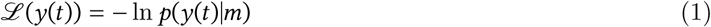

In the context of a self-organizing system, these perturbations or sensations might be the induced changes-of-state that the system undergoes in response to fluctuations from the environment. Under weak ergodicity assumptions (namely, the statistics of the steady-state distribution of the system do not diverge over time), the time-averaged surprise is the Shannon entropy of the system’s ‘sensations’:

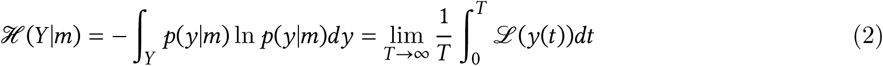

This equivalence underlies the notion that systems tend to resist dissipative forces by limiting the entropy induced by impinging fluctuations. This means such systems minimise the time-integral of surprise. To tie this dynamical formulation of surprise to the Free Energy Principle and its consequences for perception and action, we have to consider the mathematics involved in computing surprise.

### 5.2 Approximate inference via Variational Bayes

The goal of Bayesian inference is to obtain the most likely explanation for data - this means maximizing the probability of a set of parameters *x* (the causal variables or explanations), given some observations *õ*, where the tilde ∼ notation indicates a sequence of such observations over time *õ* = [*o*_1_, *o*_2_, …*o*_*T*_]^*T*^. Observations in this context are the analog of the ‘sensory fluctuations’ discussed in the previous section. The most likely explanation for observations is known as the posterior probability of parameters *x*, given data *õ*. The formula for the exact posterior is given by Bayes rule:

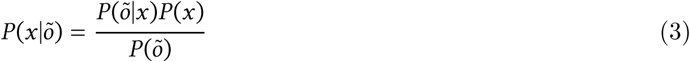

Estimating this quantity requires computing the marginal probability of data *P* (*õ*)^1^:

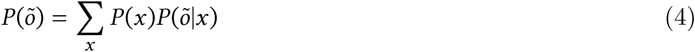

Solving this summation (in the continuous case, integration) can easily become computationally intractable for high-dimensional models, since the evidence needs to be calculated for every possible combination of parameters *x*, a computation known as ‘marginalization.’ Crucially, surprise is equal to the negative log probability of observations, as defined in Equation (1). Therefore, minimizing surprise corresponds to maximizing the probability of data, also known as (log) model evidence. This is what allows us to connect the entropy-resisting dynamics of complex systems with Bayesian inference. The marginalization in Equation (4) renders exact Bayesian inference expensive or impossible in many cases, motivating approximate inference methods. One of the leading classes of methods for approximate inference are the variational methods (Blei, Kucukelbir, & McAuliffe, 2017). Variational inference circumvents the issue of exact inference by introducing an arbitrary distribution *Q*(*x*) to replace the true posterior. This replacement is often referred to as the *variational* or *approximate posterior*. By constraining the form of the variational distribution, tractable schemes exist to optimise it in a way that (approximately) maximises evidence (Beal, 2004). This optimization occurs with respect to a quantity called the *variational free energy*, which is a computable upper-bound on surprise. The relationship between surprise and free energy can be shown as follows using the definition of joint probability distributions and Jensen’s inequality:

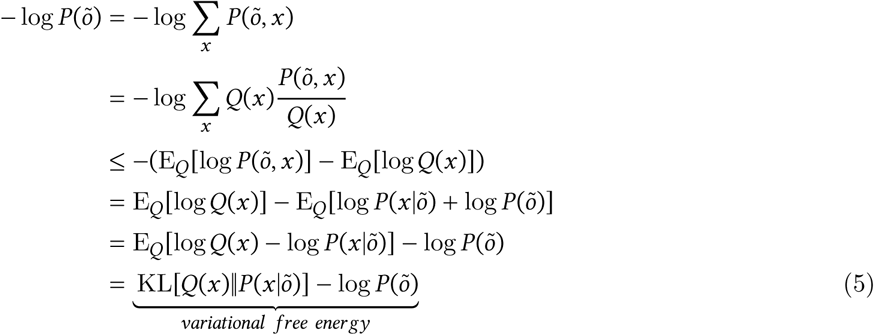

where −log*P*(*õ*) is surprise or negative log model evidence, *P* (*õ, x*) is the joint probability of observations and hidden causes, and *P* (*x*|*õ*) is the true posterior. From the last line in Equation (5) we see the free energy becomes a better approximation to surprise the more closely the variational distribution *Q*(*x*) resembles the ture posterior *P* (*x*|*õ*). This is because the first term (the Kullback-Leibler divergence^2^ between *Q*(*x*) and *P* (*x*|*õ*)) is zero when the distributions are the same. One can consider variational inference as the conversion of an integration problem (computing the marginal likelihood of observations as in Equation (4)) into an optimization problem, wherein the parameters of the variational distribution are changed to minimise *F*:

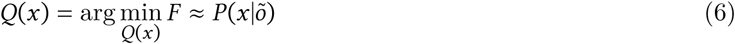

### 5.3 Active Inference and Expected Free Energy

Having discussed the variational approximation to Bayesian inference via free energy minimization, we now turn our attention to active inference. Active inference is a framework for modeling and understanding adaptive agents, premised on the idea that agents engage in approximate Bayesian inference with respect to an internal generative model of sensory data. Crucially, under active inference both action and perception are realisations of the single drive to minimise surprise. By using variational Bayesian inference to achieve this, an active inference agent generates Bayes-optimal beliefs about sources of variation in its environment by free-energy-driven optimization of an approximate posterior *Q*(*x*). This can be analogised to the idea of perception as inference, wherein perception constitutes optimizing the parameters an approximate posterior distribution over hidden states 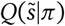, under a particular policy *π*.^3^ In the context of neural systems, it is theorised that the parameters of these posterior beliefs about states are encoded by distributed neural activity in the agent’s brain (Friston, 2008; Friston & Kiebel, 2009; Huang & Rao, 2011; Bastos et al., 2012; Parr & Friston, 2018c). Parameters of the generative model itself (such as the likelihood mapping *P* (*o*|*s*)) are hypothesised to be encoded by the network architectures, synaptic strengths, and neuromodulatory elements of the nervous system (Bogacz, 2017; Parr, Benrimoh, Vincent, & Friston, 2018; Parr, Markovic, Kiebel, & Friston, 2019).

*Action* can also be framed as a consequence of variational Bayesian inference. Under active inference, policies (sequences of actions) correspond to sequences of ‘control states’ - a type of hidden state that agents can directly influence. Actions are treated as samples from posterior beliefs about policies (Friston, Samothrakis, & Montague, 2012). However, optimizing beliefs about policies introduces an additional complication. Optimal beliefs about hidden states 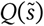 are a function of current and past observations. However, as the instantaneous free energy is a direct function of observations, it is not immediately clear how to optimise beliefs about policies when observations from the future are not available. This motivates the introduction of the *expected free energy*, or beliefs about the free energy expected at future times when pursuing a policy *π*. The free energy expected at future time point *τ* under a policy *π* is given by *G*(*π, τ*). Replacing the expectation over hidden states and outcomes in Equation 5 with the expectation over hidden states and outcomes in the future, we have:

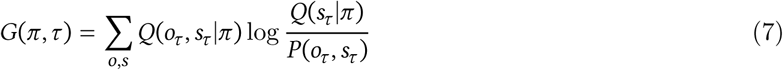

Here, we equip the agent with the prior belief that its policies minimise the free energy expected (under their pursuit) in the future.

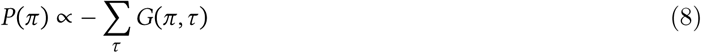

We will not derive the self-consistency of the prior belief that agents (believe they) will choose free-energy-minimizing policies, nor the full derivation of the expected free energy here. Interested readers can find the full derivations in (Friston et al., 2015; Friston, FitzGerald, et al., 2017) and (Parr & Friston, 2019). However, it is worth emphasizing that different components of the expected free energy clarify its implications for optimal behaviour in active inference agents. These components are formally related to other discussions of adaptive behaviour, such as the trade-off between exploration and exploitation. Factorizing the joint distribution *P*(*o*_*τ*_, *s*_*τ*_) in the denominator of Equation (7), we can recapitulate the expected free energy for a given time-point *τ* and policy *π* as a bound on the sum of two expectations:

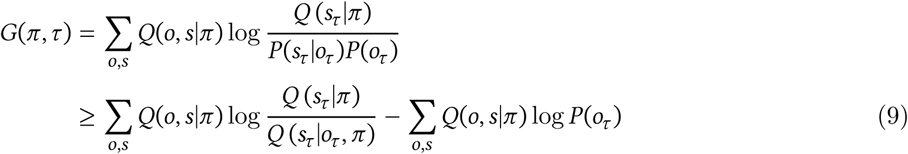

From this decomposition of the quantity bounded by the expected free energy we clarify the contribution of different kinds of ‘value’ to behaviour in active inference (Friston et al., 2013, 2015; Parr & Friston, 2017b; Mirza et al., 2018). The left term of the bottom line of Equation (9) is a term that has been called *negative information gain*. Since the most likely policies are those that *minimise* the expected free energy of their sensory consequences, this term promotes policies that disclose information about the environment by reducing uncertainty about the causes of observations. The right part of the equation can be called negative extrinsic (or instrumental) value, and minimizing this term promotes policies that result in observations that match the agent’s prior beliefs about observations, where these prior beliefs about observations will be clearer when we discuss instrumental value later on. Expected free energy is evaluated for each future timestep *τ* = *t* + 1…*T*, where *t* corresponds to the current time step and is the final time step at which the expected free energy under a policy is evaluated. Integrating over time and using the definition of expectations and the KL-divergence yields the following useful decomposition of the expected free energy **G** () of a policy:

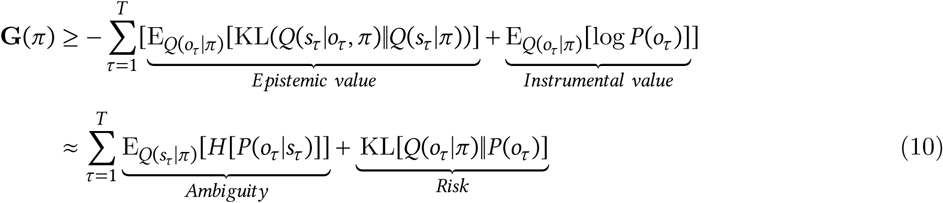

When reformulated as such, the leftmost term of the summand in the first line of Equation (10) we hereafter refer to as ‘epistemic value’ (Friston et al., 2015; Mirza et al., 2016). It is equivalent to expected Bayesian surprise or expected information gain in other accounts of information-seeking behaviour and curiosity (Linsker, 1990; Schmidhuber, 1991; Itti & Baldi, 2009; Gottlieb & Oudeyer, 2018). Such an epistemic drive has the effect of promoting actions that uncover information about hidden states via sampling informative observations. This intrinsic drive to uncover information, and its natural emergence via the minimization of expected free energy, is integral to accounts of exploratory behaviour, curiosity, salience, and related active-sensing phenomena under active inference (FitzGerald, Dolan, & Friston, 2015; Friston, Rosch, Parr, Price, & Bowman, 2017; Friston, Lin, et al., 2017; Parr & Friston, 2017b, 2018b; Mirza, Adams, Parr, & Friston, 2019). An alternative formulation of the expected free energy is given in the second line of Equation (10), where minimizing expected free energy promotes policies that reduce ‘ambiguity,’ defined as the expected uncertainty of observations, given the states expected under a policy. These notions of information gain and expected uncertainty will serve as a useful construct in understanding the behaviour of active inference agents performing hierarchical evidence accumulation and scene construction.

In order to understand how minimizing expected free energy **G** relates to the pursuit of preference-related goals or drives, we now turn to the right side of the first line of Equation (10). In order to enable instrumental or ‘non-epistemic’ goals to drive action, we supplement the agent’s generative model with an unconditional distribution over observations *P*(*o*) (sometimes called *P*(*o*|*m*), where indicates conditioning on the generative model of the agent) - this also appears in the denominator of the first line of Equation (9). By fixing certain outcomes to have high (or low) probabilities as prior beliefs, minimizing **G** imbues action selection with an apparent instrumental or exploitative component, measured by how closely observations expected under a policy align with baseline expectations. Said differently: active inference agents pursue policies that result in outcomes that they *a priori* expect to encounter. The distribution *P*(*o*) is therefore also often called ‘prior preferences.’ Encoding preferences or desires as beliefs about future sensory outcomes underwrites the known duality between inference and optimal control (Todorov, 2008; Friston, Daunizeau, & Kiebel, 2009; Friston, 2011). In the language of Expected Utility Theory (which explains behaviour by appealing to the principle of maximizing expected rewards), the logarithm of such prior beliefs is equivalent to the utility function (Zeki, Goodenough, & Zak, 2004). This component of **G** has variously been referred to as utility, extrinsic value, or instrumental value (Seth, 2015; Friston, FitzGerald, et al., 2017; Seth & Tsakiris, 2018; Biehl, Guckelsberger, Salge, Smith, & Polani, 2018); hereafter we will use the term instrumental value. A related but subtly different perspective is provided by the right-hand side of the second line of Equation (10): in this formulation, prior preferences enter the free energy through an ‘expected risk’ term. The minimization of expected risk favors actions that minimise the KL-divergence between outcomes expected under a policy and preferred outcomes, and is related to existing formulations like KL-control or risk-sensitive control (Klyubin, Polani, & Nehaniv, 2005; van den Broek, Wiegerinck, & Kappen, 2010).

### 5.4 Summary

We have seen how both perception and action emerge as consequences of free energy minimization under active inference. Perception is analogised to state estimation and corresponds to optimizing the sufficient statistics of variational beliefs over hidden states 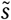. Meanwhile actions are sampled from inferred sequences of control states (policies). The likelihood of a policy is inversely proportional to the free energy *expected* under that policy. We demonstrated that expected free energy can be decomposed into the sum of two terms, which respectively encode the drive to resolve ambiguity about the hidden causes of sensory data (epistemic value) and to satisfy agent-specific preferences (instrumental value) (first line of Equation (10)). In this way active inference theoretically dissolves exploration-exploitation dilemma often discussed in decision sciences and reinforcement learning (Schmidhuber, 1991; March, 1991; Sutton & Barto, 1998) by choosing policies that minimise expected free energy. This unification of perception and action under a common Bayesian ontology underlies the power of active inference as a normative framework for studying adaptive behaviour in complex systems. In the following sections we will present a (hierarchical) Markov Decision Process model of scene construction, where stochastic motion stimuli serve as observations for an overarching scene categorization task. We then discuss perception and action under active inference in the context of hierarchical scene construction, with accompanying computational demonstrations.

## 6 Hierarchical Markov Decision Process for Scene Construction

We now introduce the hierarchical active inference model of visual foraging and scene construction. The generative model (the agent) and the generative process of the environment both take the form of a Markov Decision Process or MDP. MDPs are a simple class of probabilistic generative models where space and time are treated discretely (Puterman, 1995). In the MDP used here, states are treated as discrete samples from categorical distributions and likelihoods act as linear transformations of hidden states, mapping states at time *t* to states at time *t* + 1, or from states to observations at time *t*. This specification imbues the environment with Markovian, or ‘memoryless’ dynamics. Although the evolution of an agent’s posterior over states and actions is also constrained to obey these dynamics, the agent nevertheless violates the Markov property by constructing beliefs about the past and future with (theoretically infinite) temporal depth.

A generative model is simply a joint probability distribution over sensory observations and their latent states 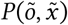, and is often factorised into the product of a likelihood and a set of marginal distributions over latent variables and hyperparameters. The discrete MDP constrains these distributions to have a particular form; here, the priors over initial states, transition and likelihood matrices are encoded as categorical distribution over a discrete set of states and samples. Agents can only directly observe sensory outcomes *õ*, making the current formulation a *partially-observed* MDP (or POMDP). Namely, the agent does not have access to true hidden states 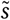, but performs inference over them by inverting the generative model to estimate the most likely sources of observations *õ*. Hierarchical models take this a step further by adding multiple layers of hidden state inference, fixing the beliefs about hidden states 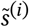 of one level to act as inferred observations *õ*^(*i*+1)^ for the level above, with associated priors and likelihoods operating at all levels. Note that as with *õ*, we use 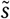 to denote a sequence of hidden states over time 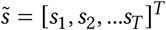

### 6.1 Hierarchical MDPs

Figure 4 summarises the structure of a generic two-layer hierarchical POMDP model, outlining relationships between random variables via a Bayesian graph and their (factorised, categorical) forms in the left panel. In the left panel of Figure 4, *õ* and 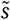 indicate sequences of observations and states over time. In the MDP model, the probability distributions that involve these sequences are expressed in a factorised fashion. The model’s beliefs about how hidden states 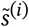 cause observations *õ*^*(i)*^ are expressed the multidimensional arrays in the the likelihood matrix **A**^(*i*),*m*^, where *i* indicates the index of the hierarchical level and *m* indicates a particular modality (Mirza et al., 2016; Friston, Rosch, et al., 2017). The (*x, y*) entry of a likelihood matrix **A**^(*i*),*m*^ prescribes the probability of observing the outcome *x* under the modality *m* at level *i*, given hidden state *y.* In this way, the columns of the **A** matrices are conditional categorical distributions over outcomes, given the hidden state indexed by the column. The dynamics that describe how hidden states at a given level *s*^(*i*)^evolve over time are given by Markov transition matrices **B** ^(*i*),*n*^ (*u*) which express how likely the next state is given the current state. Here *n* indexes a particular factor of level *i*’s hidden states, and *u* indexes a particular control state or action. Actions in this scheme are thus treated as controlled transitions between hidden states. We assume that the posterior distribution over different dimensions of hidden states factorise, leading to conditional independence between separate hidden state factors. This is known as the *mean-field assumption*, and permits efficient algorithms for updating the sufficient statistics of posterior beliefs about different hidden states (Feynman, 1998). This results in a set of relatively simple update equations for posterior beliefs and is also consistent with known features of neuroanatomy, e.g. functional segregation in the brain (Felleman & Van, 1991; Ungerleider & Haxby, 1994; Friston & Buzsáki, 2016; Mirza et al., 2016; Parr & Friston, 2018a). The hierarchical MDP formulation notably permits a segregation of timescales across layers and an according mean-field assumption on their respective free energies, such that multiple time steps of belief-updating at one level can unfold within a single time step of inference at the level above. In this way, low-level beliefs about hidden states (and policies) can be accumulated over time at a lower layer, at which point the final posterior estimate about hidden states is passed ‘up’ as an *inferred* outcome to the layer above. Subsequent layers proceed at their own characteristic (slower) timescales (Friston, Rosch, et al., 2017) to update beliefs about hidden states at their respective levels.

**Figure 4:**
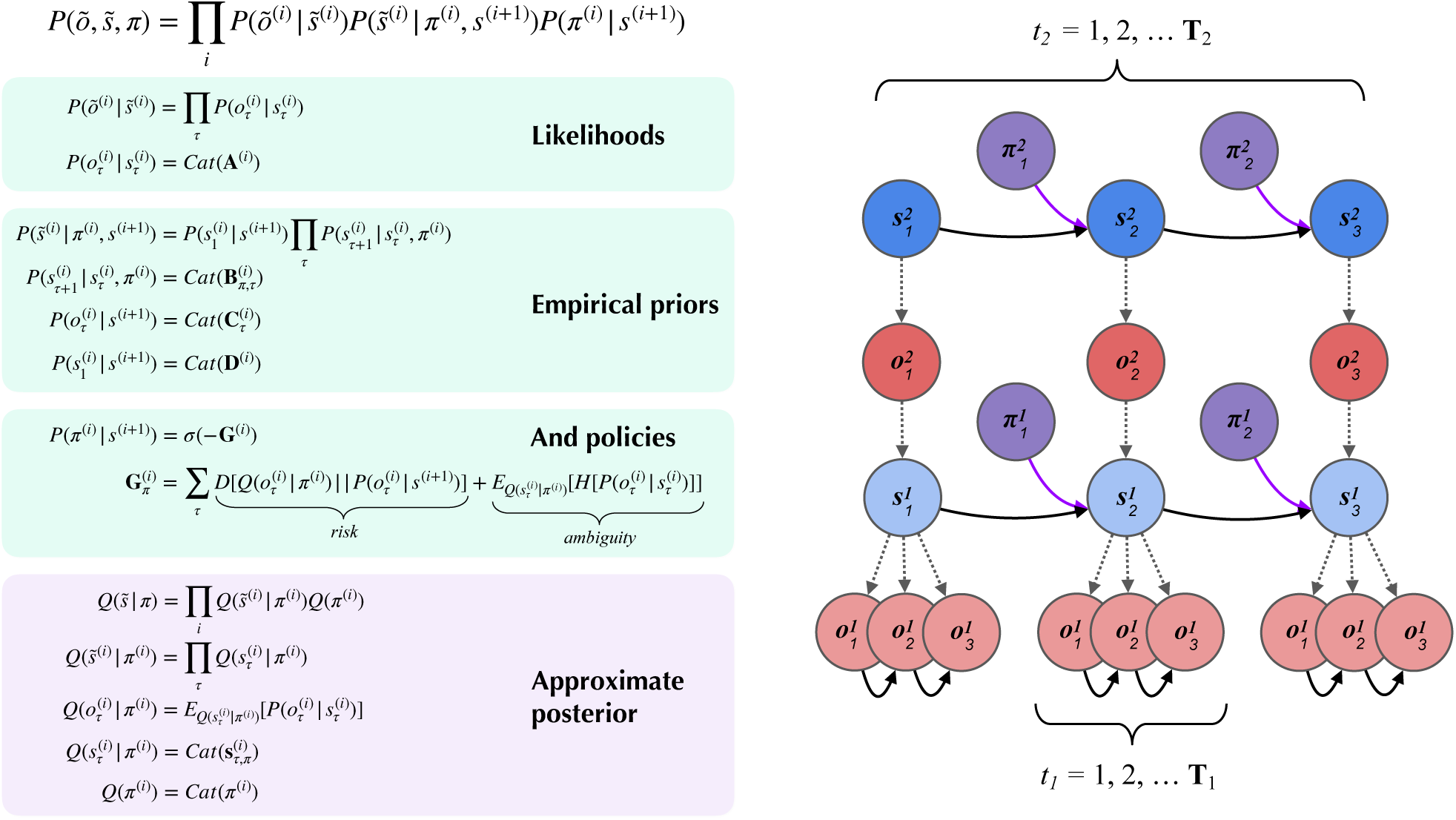
A partially-observed Markov Decision Process with two hierarchical layers. Schematic overview of a hierarchical partially-observed Markov Decision Process for both the generative process and generative model in the scene construction task. The generic forms of the likelihoods, priors, and posteriors at hierarchical levels are provided in the left panels, adapted with permission from (Friston, Rosch, et al., 2017). *Cat*(x) indicates a categorical distribution. The posterior beliefs about policies are given by a softmax function of the expected free energy of policies at a given level. The approximate (variational) beliefs over hidden states are represented via a mean-field approximation of the full posterior, such that hidden states can be encoded as the product of marginal distributions. Factorization of the posterior is assumed across hierarchical layers, across hidden state factors (see the text and Figures 6 - 7 for details on the meanings of different factors), and across time.

Figure 5 gives an overview of the belief update equations for state inference (hidden states 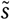) and policy evaluation (posterior over policies) under active inference. Indices here only refer to policy and time step (for the sake of clarity we don’t include the empirical priors from other hierarchical levels). Note that here, instead of directly evaluating the solution for states with lowest free energy **s**\*, we use a marginal message passing routine to perform a gradient descent on the variational free energy at each time step, where posterior beliefs about hidden states and policies are incremented using prediction errors (see Figure 5 legend for more details). In the context of deep temporal models, these equations proceed independently at each level of the hierarchy at each time step. At lower levels the posterior over states at the first timestep 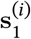 is an initialised as the expected observations **o**^(*i*+1)^ from the level above, and inferred observations at higher levels are inherited as the final posterior beliefs 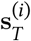 from lower levels. This update scheme may sound complicated; however, when expressed in terms of expected states, it reduces to a remarkably simple scheme that, crucially, looks very much like neuronal processing: see (Friston et al., 2015) for details. The right side of Figure 5 shows a simple schematic of how particular random variables that make up generative model may map onto defined neuroanatomical areas, within which the belief-updating proceeds according to the equations in Panels **A** - **C**. Evidence for the sort of hierarchical processing entailed by such generative models abounds in the brain, and is the subject of a wealth of empirical and theoretical neuroscience research (Lee & Mumford, 2003; Hasson, Yang, Vallines, Heeger, & Rubin, 2008; Friston, 2008; Friston, Parr, & de Vries, 2017; Runyan, Piasini, Panzeri, & Harvey, 2017; Pezzulo, Rigoli, & Friston, 2018).

**Figure 5:**
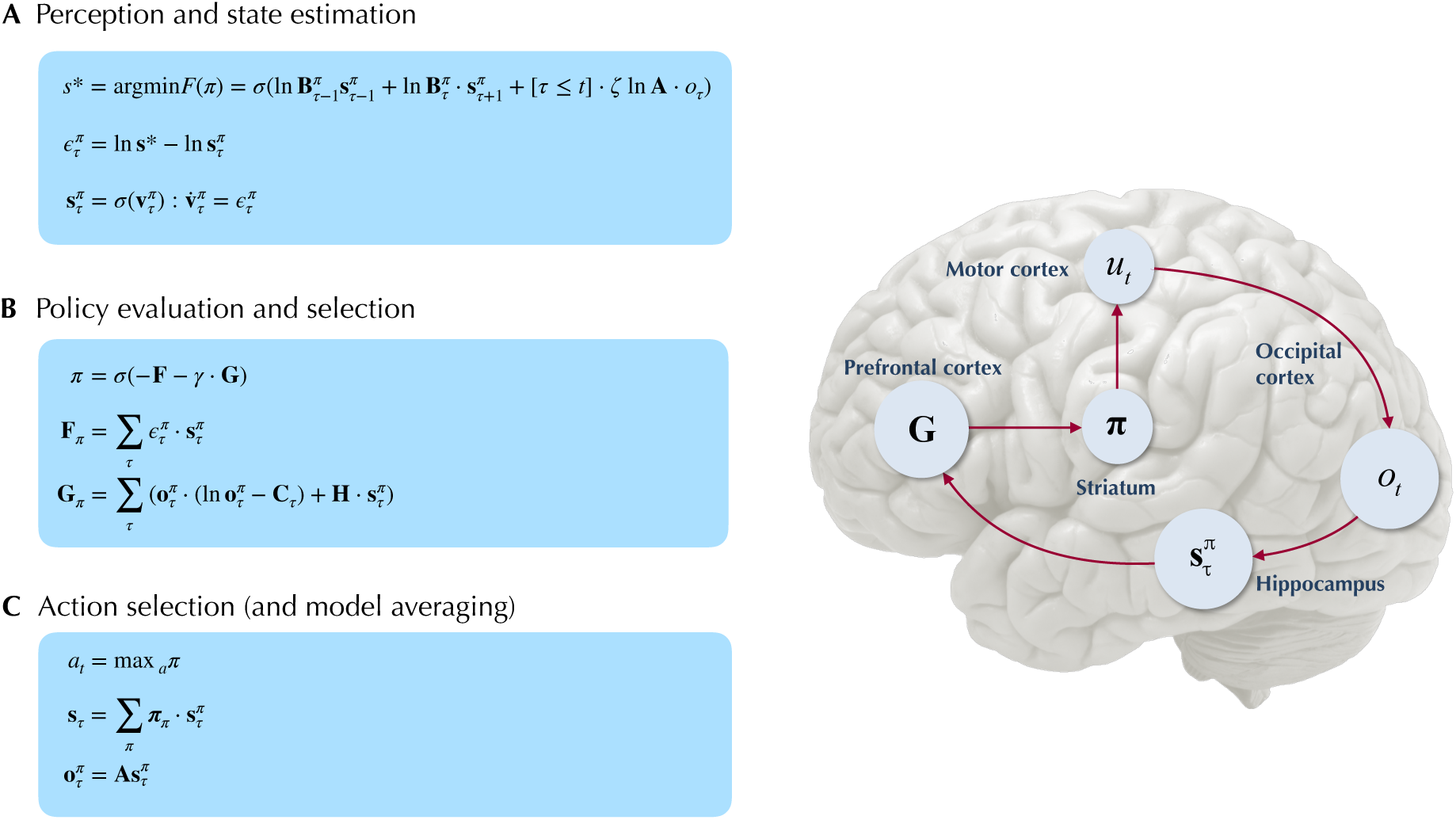
Belief-updating under active inference. Overview of the update equations for posterior beliefs under active inference. (**A**) shows the optimal solution for posterior beliefs about hidden states **s**^*^ that minimises the variational free energy of observations. In practice the variational posterior over states is computed as a marginal message passing routine (Parr, Markovic, et al., 2019), where prediction errors 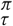 minimised over time until some criterion of convergence is reached (≈ 0). The prediction errors measure the difference between the current log posterior over states ln 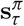 and the optimal solution ln **s**^**\***^. (**B**) shows how posterior beliefs about policies are a function of the free energy of states expected under policies **F** and the expected free energy of policies **G. F** is a function of state prediction errors and expected states, and **G** is the expected free energy of observations under policies, shown here decomposed into the KL divergence between expected and preferred observations or risk 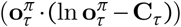 and the expected entropy or ambiguity 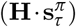. A precision parameter *γ scales* the expected free energy and serves as an inverse temperature parameter for a softmax normalization *σ* of policies. (**C**) shows how actions are sampled from the posterior over policies, and the posterior over states is updated via a Bayesian model average, where expected states are averaged under beliefs about policies. Finally, expected observations are computed by passing expected states through the likelihood of the generative model. The right side shows a plausible correspondence between several key variables in an MDP generative model and known neuroanatomy. For simplicity, a hierarchical generative model is not shown here, but one can easily imagine a hierarchy of state inference that characterises the recurrent message passing between lower-level occipital areas (e.g., primary visual cortex) through higher level visual cortical areas, and terminating in ‘high-level,’ prospective and policy-conditioned state estimation in areas like the hippocampus.

### 6.2 From Motion Discrimination to Scene Construction: a Nested Inference Problem

We now introduce the deep, temporal model of scene construction using the task discussed in Section 4 as our example (Figure 6). We formulate perception and action with a hierarchical POMDP consisting of two distinct layers that are solved via active inference. The first, shallowest level (Level 1) is an MDP that updates posterior beliefs about the most likely cause of visual stimulation (RDM direction), where we model the ongoing contents of single fixations - the stationary periods of relative retinal-stability between saccadic eye movements. This inference is achieved with respect to the (spatially-local) visual stimuli underlying current foveal observations. A binary policy is also implemented, encoding the option to continue holding fixation (and thus keep sampling the current stimulus) or to interrupt sampling and terminate updates at the lower level. The second, higher level (Level 2) is another MDP that performs inference at a slower timescale, with respect to the overarching hidden scene that describes the current trial. Here, we enable policies that realise visual foraging. These policies encode controlled transitions between different states of the oculomotor system, serving as a model of saccadic eye movements to different parts of the visual array. This way of encoding saccades is inspired by the original scene construction formulation in (Mirza et al., 2016). We will now discuss both layers individually and translate different elements of the MDP generative model and environment to task-relevant parameters and the beliefs of the agent.

**Figure 6:**
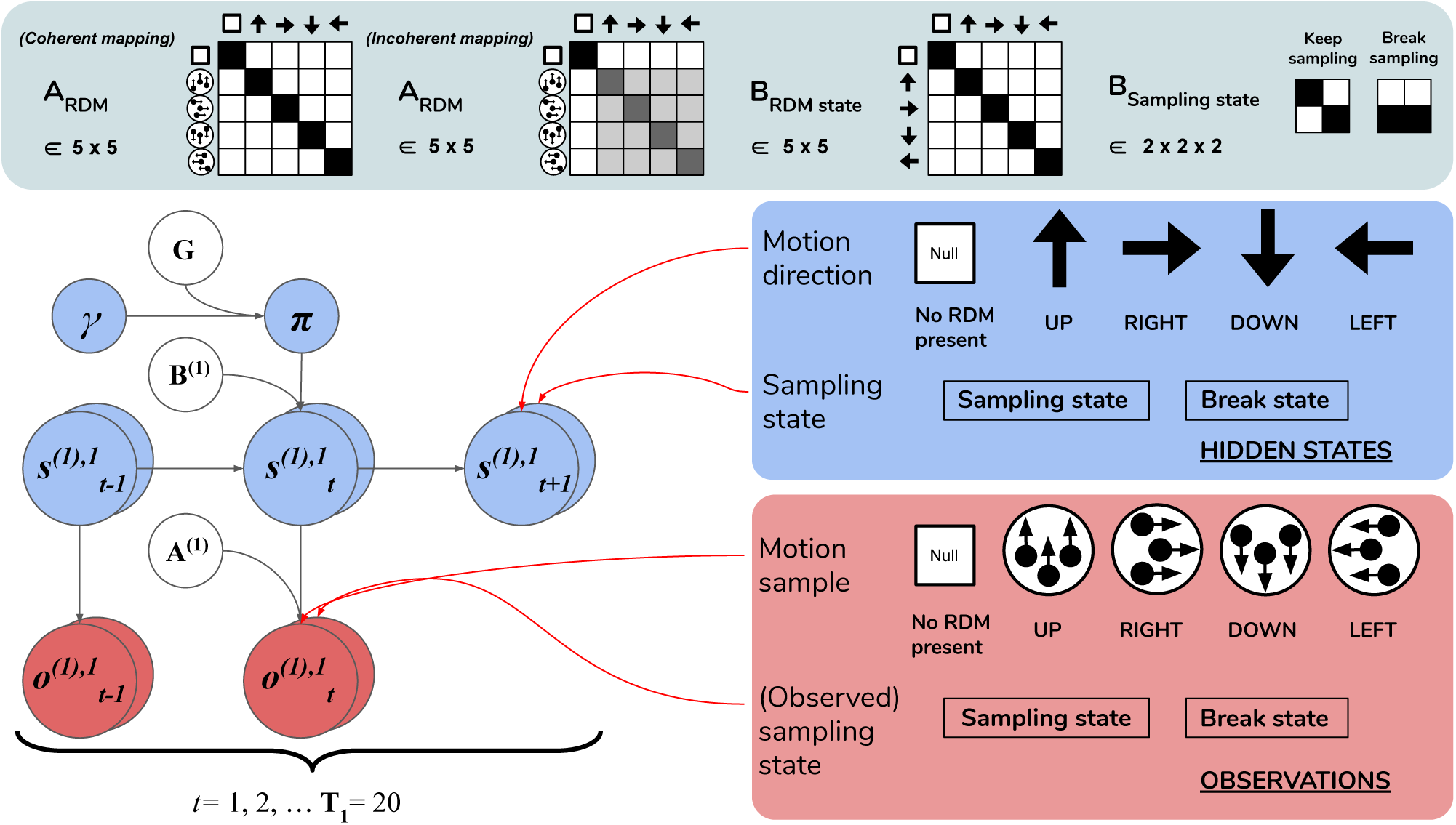
Level 1 MDP. Schematic of the graphical model and associated likelihoods for **Level 1** of the hierarchical generative model and process in scene construction (see Section 6.2.1 for details). At this level hidden states correspond to: 1) the motion direction **s**^(1),1^ underlying visual observations at the currently-fixated region of the visual array and 2) the sampling state **s**^(1),1^, which can be changed via policies that select the appropriate **B** matrix-encoded state transition. The first hidden state factor **s**^(1),1^ can either correspond to a state with no motion signal (‘Null’, in the case when there is no RDM or a categorization decision is n being made) or assume one of the four discrete values corresponding to the four cardinal motion directions. At each time step of the generative process, the current hidden state **s**^1^ is probabilistically mapped to a motion observation via the first-factor likelihood **A**^(1),1^ (shown in the top panel as **A**_RDM_). The entropy of the columns of this mapping can be used to parameterise the coherence of the RDM stimulus, such that the true motion states **s**^1^ cause motion observations **o**^1^ with varying degrees of fidelity. This is demonstrated by two exemplary **A**RDM state matrices in the top panel (these correspond to **A**^(1),1^): the left-most matrix shows a noiseless, ‘coherent’ mapping, analogised to the situation of when an RDM consists of all dots moving in the same direction as described by the true hidden state; the matrix to the right of the noiseless mapping corresponds to an incoherent RDM, where instantaneous motion observations may assume directions different than the true motion direction state, with the frequency of this deviation encoded by probabilities stored in the corresponding column of **A**_RDM_. The motion direction state doesn’t change in the course of a trial (see the identity matrix shown in the top panel as **B**_RDM_, which simply maps the hidden state to itself at each subsequent time step). The second hidden state factor **s**^(1),2^ encodes the current ‘sampling state’ of the agent; there are two levels under this factor: the ‘**Keep-sampling**’ state or the ‘**Break-sampling**’ state. The break state is a sink in the transition dynamics such that once it is entered, it cannot be exited (see the right-most matrix of the **B**_Sampling state_ likelihood). Entering the ‘**Break-sampling**’ state terminates the temporal updates at Level 1. The ‘**Keep-sampling**’ state enables the continued generation of motion observations as samples from the the likelihood mapping **A**^(1),1^.**A** ^(1),2^(the ‘proprioceptive’ likelihood, not shown for clarity) deterministically maps the current sampling state **s**^(1),2^ to an observation **o**^(1),2^ (bottom row of lower right panel).

#### 6.2.1 Level 1: Motion discrimination via motion sampling over time

Lowest level (Level 1) beliefs are updated as the agent encounters a stream of ongoing, potentially ambiguous visual observations - the instantaneous contents of an individual fixation. The hidden states at this level describe a distribution over motion directions, which parameterise the true state of the random motion stimulus within the currently-fixated quadrant. Observations manifest as a sequence of stochastic motion signals that are samples from the true hidden state distribution.

The generative model has an identical form as the generative process (see above) used to generate the stream of Level 1 outcomes. Namely, it is comprised of a set of likelihoods and transitions as the dynamics describing the ‘real’ environment (Figure 6). In order to generate a motion observation, we sample the probability distribution over motion direction given the true hidden state using the Level 1 generative process likelihood matrix **A**^(1),1^. For example, if the current true hidden state at the lower level is **2** (implying that an RDM stimulus of **UP**wards motion occupies the currently fixated quadrant), stochastic motion observations are sampled from the *second* column of the generative likelihood mapping **A**^(1),1^. The precision of this column-encoded distribution over motion observations determines how often the sampled motions will be **UP**wards signals and thus consistent with the true hidden state. The entropy or ambiguity of this likelihood mapping operationalises sensory uncertainty and in this case, motion incoherence. For more details on how states and outcomes are discretised in the generative process, see Figure 6 and its legend.

Inference about the motion direction (Level 1 state estimation) roughly proceeds as follows: **1)** at time *t* a motion observation 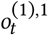 is sampled from the generative process **A**^(1),1^; **2)** posterior beliefs about the motion direction at the current timestep 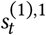 are updated using a gradient descent on the variational free energy. In addition, we included a second, controllable hidden state factor at Level 1 that we refer to as the abstract ‘sampling state’ of the agent. We include this in order to enable policies at this level, which entail transitions between the two possible values of this control state. These correspond to the choice to either keep sampling the current stimulus or break sampling. These policies are stored as two 2 x 2 transition matrices in **B**^(2),2^, where each transition matrix **B**^(2),2^(*u*)encodes the probability of transitioning to ‘**Keep-sampling**’ or ‘**Break-sampling**,’ given an action *u* and occupancy in one of the two sampling states. Selecting the first action keeps the Level 1 MDP in the ‘**Keep-sampling**’ state, triggering the generation of another motion observation from the generative process. Engaging the 2nd ‘**Break-sampling**’ policy moves the agent’s sampling regime into the second state and terminates any further updates at Level 1. At this point the latest posterior beliefs from Level 1 are sent up as observations for Level 2. It is worth noting that implementing ‘breaking’ the MDP at the lower level as an explicit policy departs from the original formulation of deep, temporal active inference. In the formulation developed in (Friston, Rosch, et al., 2017), termination of lower level MDPs occurs once the entropy of the lower-level posterior over the hidden states (only those factors that are linked with the level above) is minimised beyond a fixed value^4^. We chose to treat breaking the first level MDP as an explicit policy in order to formulate behaviour in terms of the same principles that drive action selection at the higher level - namely, the expected free energy of policies. In the Simulations section we explore how the dynamic competition between the ‘**Break**-’ and ‘**Keep-sampling**’ policies induces an unexpected distribution of break latencies.

We fixed the maximum temporal horizon of Level 1 (hereafter *T*_1_) to be 20 time steps, such that if the ‘**Break-sampling**’ policy is not engaged before *t* = 20 (implying that ‘**Keep-sampling**’ has been selected the whole time), Level 1 automatically terminates after the 20th time step and the final posterior beliefs are passed up as outcomes for Level 2.

#### 6.2.2 Level 2: Scene Inference and Saccade Selection

After beliefs about the state of the currently-foveated visual region are updated via active inference at Level 1, the resulting posterior belief about motion directions is passed up to Level 2 as a belief about observations. These observations (which can be thought of as the inferred state of the visual stimulus at the foveated area) are used to update the statistics of posterior beliefs over the hidden states operating at Level 2 (specifically, the hidden state factor that encodes the identity of the scene, e.g. **UP-RIGHT**). Hidden states at Level 2 are segregated into two factors, with corresponding posterior beliefs about them updated independently.

The first hidden state factor corresponds to the scene identity. As described in Section 4, there are four possible scenes characterizing a given trial: **UP-RIGHT, RIGHT-DOWN, DOWN-LEFT**, and **LEFT-UP**. The scene determines the identities of the two RDMs hiding throughout the four quadrants - e.g. when the scene is **UP-RIGHT**, one **UP**wards-moving RDM is found in one of the four quadrants, and a **RIGHT**wards-moving RDM is found in another quadrant. The quadrants that are occupied by RDMs for a given scene is random, meaning that agents have to forage the 2 x 2 array for the RDMs in order to infer the scene. We encode the scene identities as well as their ‘spatial permutability’ (with respect to quadrant-occupancy) by means of a single hidden state factor that exhaustively encodes the unique combinations of scenes and their spatial configurations. This first hidden state factor is thus a 48-dimensional state distribution (4 scenes x 12 possible spatial configurations - see Figure 7 for visual illustration).

**Figure 7:**
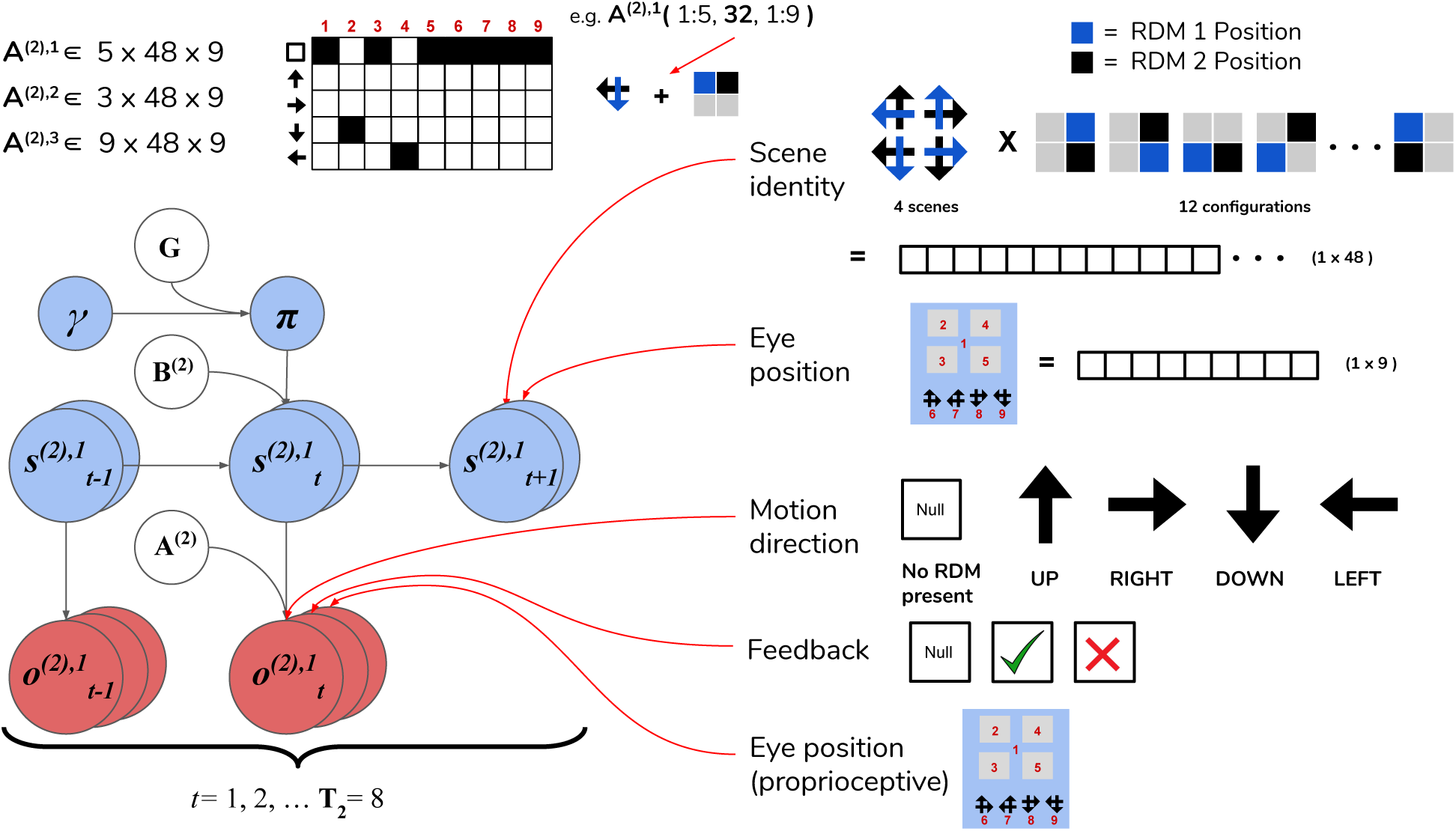
Level 2 MDP. Schematic of the MDP generative model and generative process operating at **Level 2** of hierarchical scene construction. Hidden states consist of two factors, one encoding the scene identity and another encoding the eye position (i.e., current state of the oculomotor system). The first hidden state factor encodes the scene identity of the trial in terms of two unique RDM directions occupy two of the quadrants (4 possible scenes as described in the top right panel) and spatial configuration (one of 12 unique ways to place two RDMs in four quadrants). This yields a dimensionality of 48 for this hidden state factor (4 scenes x 12 spatial configurations). The second hidden state encodes the eye position, which is initialised to be in the center of the quadrants (Location 1). The next four values of this factor index the four quadrants (2 - 5), and the last four are indexes for the choice locations (the agent fixates one of these four options to guess the scene identity). Outcomes at Level 2 are characterised by three modalities: the first modality indicates the visual stimulus (or lack thereof) at the currently-fixated location. An example likelihood matrix for this first modality is shown in the upper left, showing the conditional probabilities for visual outcomes when the 1^st^ factor hidden state has the value 32. This corresponds to the scene identity **DOWN-LEFT** under spatial configuration 8 (the RDMs occupy quadrants indexed as Locations 2 and 4). The last two likelihood arrays **A**^(2),2^ and **A**^(2),3^ are not shown for clarity; the **A**^(2),2^ likelihood encodes the joint probability between the trial feedback (Null, Correct, Incorrect) as a function of the current hidden scene and the location of the agent’s eyes, while A^(2),3^ is an unambiguous proprioceptive mapping that signals to the agent the location of its own eyes.

The second hidden state factor corresponds to the current spatial position that’s being visually fixated - this can be thought of as a hidden state encoding the current configuration of the agent’s eyes. This hidden state factor has 9 possible states: the first state corresponds to an initial position for the eyes (i.e. a fixation region in the center of the array); the next four states (indices 2 - 5) correspond to the fixation positions of the four quadrants in the array, and the final four states (6 - 9) correspond to categorization choices (i.e. a saccade which reports the agent’s guess about the scene identity). The states of the first and second hidden state factors jointly determine which observation is sampled at each timestep on Level 2.

Observations at this level comprise 3 modalities. The first modality encodes the identity of the visual stimulus at the fixated location and is identical in form to the first hidden state factor at Level 1: namely, it can be either the ‘Null’ outcome (when there is no visual stimulus at the fixated location) or one of the four motion directions. The likelihood matrix for the first-modality on Level 2, namely A^(2),1^, consists of probabilistic mappings from the scene identity /spatial configuration (encoded by the first hidden state factor) and the current fixation location (the second hidden state factor) to the stimulus identity at the fixated location - e.g. if the scene is **UP-RIGHT** under the configuration where the **UP**wards-moving RDM is in the upper left quadrant and the **RIGHT**wards-moving RDM is in the upper right quadrant and the current fixation location (the second hidden state) is the upper left quadrant, then the likelihood function will determine the first-modality observation at Level 2 to be **UP**. When the agent is fixating either an empty quadrant, the starting fixation location, or one of the response options (locations 6 - 9), the observation in the first modality is **Null**. The likelihood functions are deterministic and identical in both the generative model and generative process - this imbues the agent with a kind of ‘prior knowledge’ of the (deterministic) mapping between the scenes and their respective visual manifestations in the 2 x 2 grid. The second observation modality is a ternary variable that returns feedback to the agent based on its scene categorization performance - it can assume the values of ‘No Feedback,’ ‘Correct,’ or ‘Incorrect.’ Including this observation modality (and prior beliefs about the relative probability of its different values) allows us to endow agents with the drive to report their guess about the scene, and to do so accurately in order to maximise the chance of receiving correct feedback. The likelihood mapping for this modality **A**^(2),2^is structured to return a ‘No Feedback’ outcome in this modality when the agent fixates any area besides the response options, and returns ‘Correct’ or ‘Incorrect’ once the agent makes a saccade to one of the response options (locations 6 - 9) - the particular value it takes depends jointly on the true hidden scene and the scene identity that the agent has guessed. We will further discuss how a drive to respond accurately emerges when we describe the prior beliefs parameterised by the **C** and **D** arrays. The final observation modality at Level 2 is a proprioceptive mapping (similar to ‘sampling-state’ modality at Level 1) that unambiguously signals which location the agent is currently fixating via a 9 x 9 identity matrix **A**^(2),3^.

The transition matrices at Level 2, namely **B**^(2),1^ and **B**^(2),2^, describe the dynamics of the scene identity and of the agent’s oculomotor system, respectively. We assume the dynamics that describe the scene identity are both uncontrolled and unchanging, and thus fix **B**^(2),1^ to be an identity matrix that ensures the scene identity/spatial configuration is stable over time. As in earlier formulations (Friston, Adams, et al., 2012; Mirza et al., 2016; Mirza, Adams, Friston, & Parr, 2019) we model saccadic eye movements as transitions between control states in the 2nd hidden state factor. The dynamics describing the eye movement from the current location to a new location is encoded by the transition array **B**^(2),2^ (e.g. if the action taken is 3 then the saccade destination is described by a transition matrix that contains a row of 1s on the third row, mapping from any previous location to location 3).

Inference and action selection at Level 2 proceeds as follows: based on the current hidden state distribution and Level 1’s likelihood mapping **A**^(1),1^ (the generative process), observations are sampled from the three modalities. The observation under the first-modality at this level (either ‘Null’ or a motion direction parameterizing an RDM stimulus) is passed down to Level 1 as the initial true hidden state. The agent also generates expectations about the first-modality observations via **A**^(1),1^ · *s*_*t*_, where **A**^(1),1^ is the generative model’s likelihood and *st* is the latest posterior density over hidden states (factorised into scene identity and fixation location). This predictive density over (first-modality) outcomes serves as an empirical prior for the agent’s beliefs about the hidden states in the first factor - motion direction - at Level 1. Belief-updating and policy selection at Level 1 then proceeds via active inference using the empirical priors inherited from Level 2 in addition to its own generative model and process (as described in Section 6.2.1). Once the motion observations and belief updating terminates at Level 1, the final posterior beliefs about the 1st factor hidden states are passed to Level 2 as inferred observations of the first modality. The belief updating at Level 2 proceeds as usual, where observations are integrated using Level 2’s generative model to form posterior beliefs about hidden states and policies that minimise free energy. An action is then sampled from the policy posterior (which is a softmax function of negative expected free energy under all policies), which changes hidden states in the next time step and generate a new observation, thus closing the action-perception cycle. In this spatiotemporally ‘deep’ version of scene construction, we see how a temporally-extended process of active inference at the lower level (capped at **T**_1_ = 20 time steps in our case) can be nested within a single time step of a higher-level process, endowing such generative models with a flexible, modular form of temporal depth. Also note the asymmetry in informational scheduling across layers, with posterior beliefs about those hidden states linked with the higher level being passed up as evidence for outcomes at the higher level, with observations at the higher level being passed down as empirical priors over hidden states at the lower level.

#### 6.2.3 Priors

In addition to the likelihood **A** and **B** arrays that prescribe the probabilistic relationships between variables at each level, the generative model is also equipped with prior beliefs over observations and hidden states that are respectively encoded in the so-called **C** and **D** arrays. See Figure 8 for schematic analogies for these arrays and their elements for the two hierarchical levels.

**Figure 8:**
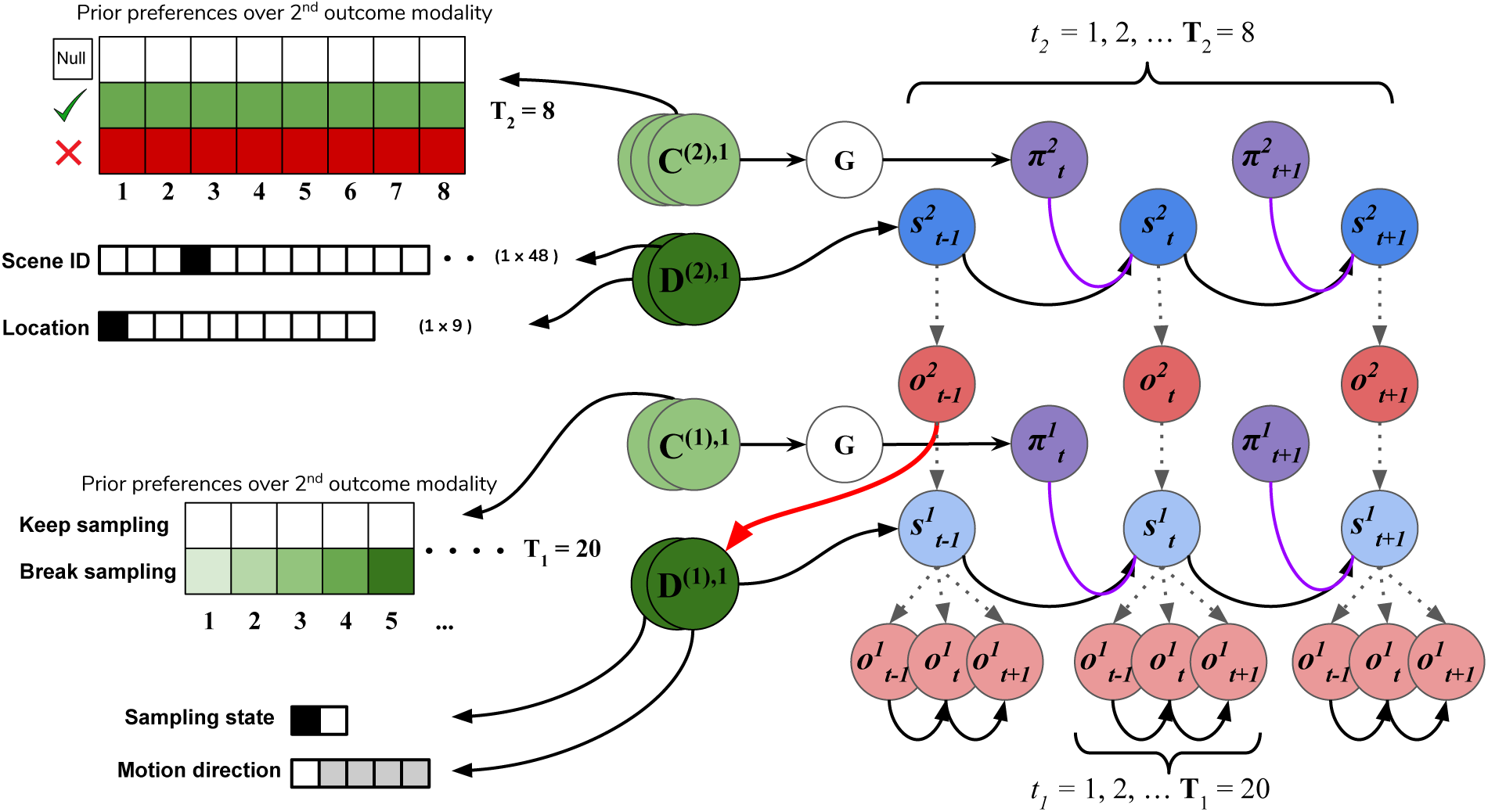
C’s and D’s. Prior beliefs over observations and hidden states for both hierarchical levels. Note that superscripts here index the hierarchical level, and separate modalities/factors are indicated by overlapping circles and with text. At the highest level (Level 2), prior beliefs about second-modality outcomes (**C**^(2),2^) encode the agent’s beliefs about receiving correct and avoiding incorrect feedback. Prior beliefs over the other outcome modalities (**C**^(2),1^ and **C**^(2),3^) are all trivially zero. These beliefs are stationary over time and affect saccade selection at Level 2 via the expected free energy of policies **G**. Prior beliefs about hidden states **D**^(2)^ at this level encode the agent’s initial beliefs about the scene identity and the location of their eyes. This prior over hidden states can be manipulated to put the agent’s beliefs about the world at odds with the actual hidden state of the world. At Level 1, the agent’s preferences about being in the ‘**Break-sampling**’ state increases over time and is encoded in the preferences about second modality outcomes (**C**^(1),2^), which corresponds to the agents umambiguous perception of its own sampling state. Finally, the prior beliefs about initial states at Level 1 (**D**^1^) correspond to the motion direction hidden state (the RDM identity) as well as which sampling-state the agent is in. Crucially, the first factor of these prior beliefs **D**^(1),1^ is defined as the expected Level 2 observations about motion direction (the first modality) *Q*(*o*^(2),1^|*s*^(2)^) (see the red arrow). The prior over hidden states at Level 1 is thus called an empirical prior as it is inherited from Level 2. The first factor of the true hidden state at Level 1 *s*^(1),1^ is the sampled discrete observation from the Level 2 likelihood matrix in the generative process.

The **C** array contains what are often called the agent’s ‘preferences’ *P*(*o*) and encodes the agent’s prior beliefs about observations (an unconditional probability distribution). Rather than an explicit component of the generative model, the prior over outcomes is absorbed into the prior over policies *P*(*π*), which is described in Section 5.3. Policies that are more likely to yield observations that are deemed probable under the prior (expressed in terms of agent’s preferences *P*(*o*)) will have less expected free energy and thus be more likely to be chosen. *Instrumental value* or expected utility measures the degree to which the observations expected under a policy correlate with prior beliefs about those observations. For categorical distributions, evaluating instrumental value amounts to taking the dot product of the (policy-conditioned) posterior predictive density over observations *Q*(*o*_*τ*_|*π*) with the log probability density over outcomes log. This reinterpretation of preferences as prior beliefs about observations allows us to discard the classical notion of a ‘utility function’ as postulated in fields like reward neuroscience and economics, instead explaining both epistemic and instrumental behaviour using the common currency of log-probabilities and surprise. In order to motivate agents to categorise the scene, we embed a self-expectation of accuracy into the **C** array of Level 2; this manifests as a high prior expectation of receiving ‘Correct’ feedback (a relative log probability of +2 nats) and an expectation that receiving ‘Incorrect’ feedback is unlikely (relative log probability of -4 nats). The remaining outcomes of the other modalities at Level 2 have equal log-probability in the agent’s prior preferences, thus contributing identically (and uninformatively) to instrumental value. At Level 1 we encoded a form of ‘urgency’ using the **C** matrix; we encoded the prior belief that the probability of observing the ‘**Break-sampling**’ state (via the umambiguous mapping *A*^(1),2^) increases over time. This necessitates that the complementary probability of remaining in the ‘**Keep-sampling**’ state decreases over time. Equipping the Level 1 MDP with such preferences generates a tension between the epistemic drive to resolve uncertainty about the hidden state of the currently-fixated stimulus and the ever-strengthening prior preference to terminate sampling at Level 1. In the simulation results to follow, we explore this tension more explicitly and report an interesting yet unexpected relationship between sensory uncertainty and fixational dwell time, based on the dynamics of various contributions to expected free energy.

Finally, the **D** array encodes the agent’s initial (prior) beliefs over hidden states in the environment. By changing prior beliefs about the initial states, we can manipulate an agent’s beliefs about the environment independently of the true hidden states characterizing that environment. In the section 7.2 below we describe the way we parameterise the first hidden state factor of the Level 2 **D** matrix to manipulate prior beliefs about the scene. The second hidden state factor at Level 2 (encoding the saccade location) is always initialised to start at Location 1 (the generic ‘starting’ location). At Level 1, first-factor of the **D** matrix (encoding the true motion direction of an RDM) is initialised to the posterior expectations from Level 2. The second-factor belief about initial hidden states (encoding the sampling state) is set as the ‘**Keep-sampling**’ state.

In the following sections, we present hierarchical active inference simulations of scene construction, in which we manipulate the uncertainty associated with beliefs at different levels of the generative model to see how uncertainty differentially affects inference across levels in uncertain environments.

## 7 Simulations

Having introduced the hierarchical generative model for our RDM-based scene construction task, we will now explore behaviour and belief-formation in the context of hierarchical active inference. In the following sections we study different aspects of the generative model through quantitative simulations. We relate parameters of the generative model to both ‘behavioural’ read-outs (such as sampling time, categorization latency and accuracy) as well as the agents’ internal dynamics (such as the evolution of posterior beliefs, the contribution of different kinds of value to policies, etc.). We then discuss the implications of our model for studies of hierarchical inference in noisy, compositionally-structured environments.

### 7.1 Manipulating Sensory Precision

Figures 9 and 10 show examples of deep active inference agents performing the scene construction task under two levels of motion coherence (high and low, respectively for Figure 9 and Figure 10), which is equivalent to the reliability of motion observations at Level 1. In particular, we operationalise this uncertainty via an inverse temperature *p* that parameterises a softmax transformation on the columns of the Level 1 likelihood mapping to RDM observations **A**^(1),1^ Each each column of **A**^(1),1^ is initialised as a ‘one-hot’ vector that contains a probability of 1 at the motion observation index corresponding to the true motion direction, and 0s elsewhere. As *p* decreases, **A** deviates further from the identity matrix and Level 1 motion observations become more degenerate with respect to the hidden state (motion direction) underlying them. Note that this parameterization of motion incoherence only pertains to the last four rows/columns of **A** ^(1),1^, as the first row/column of the likelihood (**A**^(1),1^(1, 1)) corresponds to observations about the ‘Null’ hidden state, which is always observed unambiguously when it is present. In other words, locations that do not contain RDM stimuli are always perceived as ‘Null’ in the first modality with certainty.

**Figure 9:**
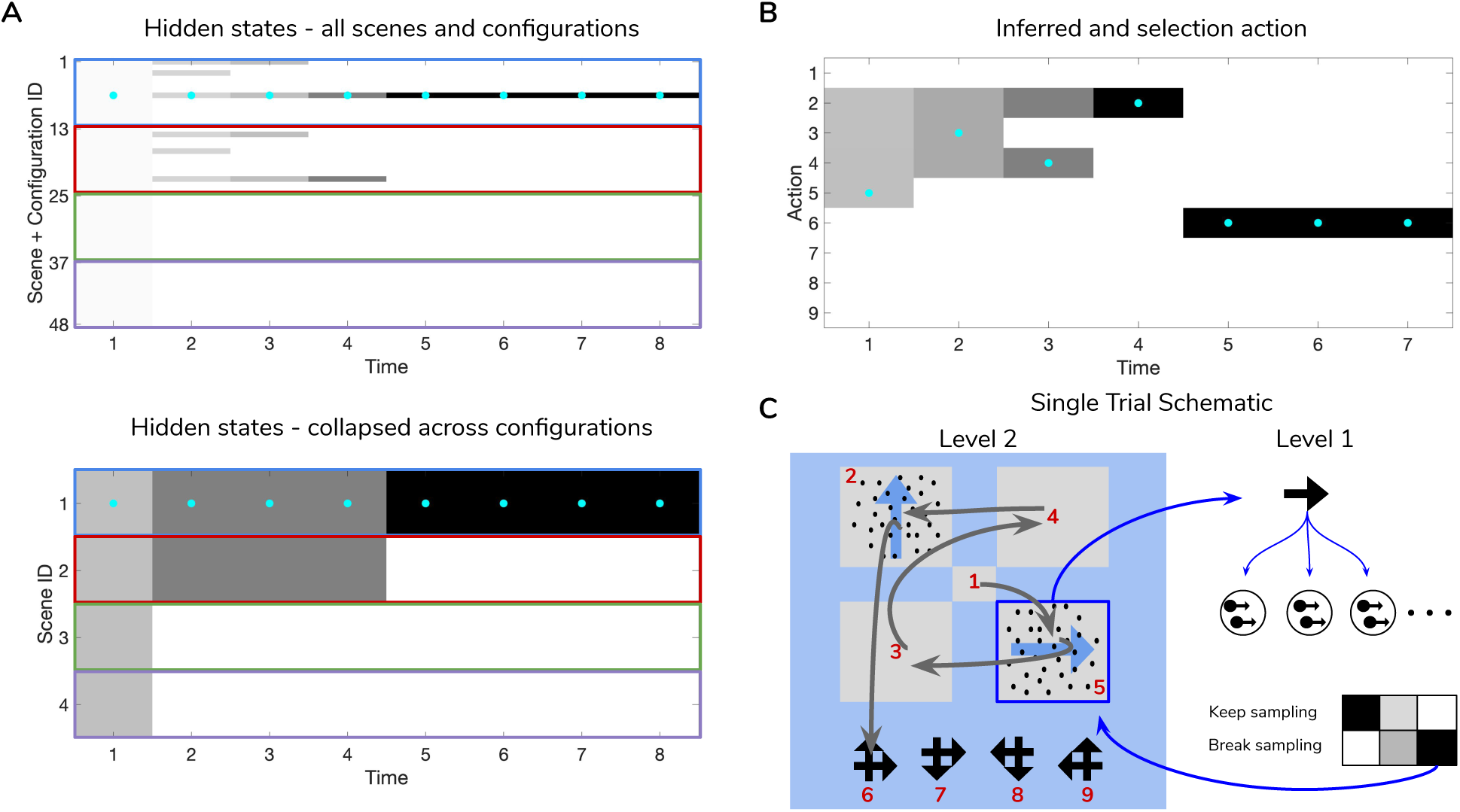
Simulated trial of scene construction under high sensory precision. (**A**): The evolution of posterior beliefs about scene identity - the first factor of hidden states at Level 2 - as a deep active inference agent explores the visual array. In this case, sensory precision at Level 1 is high, meaning that posterior beliefs about the motion direction of each RDM-containing quadrant are resolved easily, resulting in fast and accurate scene categorization. Cells are gray-scale colored according to the probability of the belief for that hidden state and time index (darker colors correspond to higher probabilities). Cyan dots indicates the true hidden state at each time step. The top row of (**A**) shows evolving beliefs about the fully-enumerated scene identity (48 possibilities), with every 12 configurations highlighted with a differently-colored bounding box, correspond to beliefs about each type of scene (i.e., **UP-RIGHT, RIGHT-DOWN, DOWN-LEFT, LEFT-UP**). The bottom panel shows the collapsed beliefs over the four scenes, computed by summing the hidden state beliefs across the 12 spatial configurations. (**B**): Evolution of posterior beliefs about actions (fixation starting location not shown), culminating in the categorization decision (here, the scene was categorised as **UP-RIGHT**, corresponding to a saccade to location 6. (**C**): Visual representation of the agent’s behaviour for this trial. Saccades are depicted as curved gray lines connecting one saccade endpoint to the next. Fixation locations (corresponding to 2^nd^ factor hidden state indices) are shown as red numbers. The Level 1 active inference process occurring within a single fixation is schematically represented on the right side, with individual motion samples shown as issued from the true motion direction via the low level likelihood **A**^(1),1^. Tha agent observes the true RDM at Level 1 and updates its posterior beliefs about this hidden state. As uncertainty about the RDM direction is resolved, the ‘**Break-sampling**’ action becomes more attractive (since epistemic value contributes increasingly less to the expected free energy of policies). In this case, the sampling process at Level 1 is terminated after only 3 timesteps, since the precision of the likelihood mapping is high (*p* = 5.0) which relates to the speed at which uncertainty is resolved about the RDM motion direction - see the text for more details.

**Figure 10:**
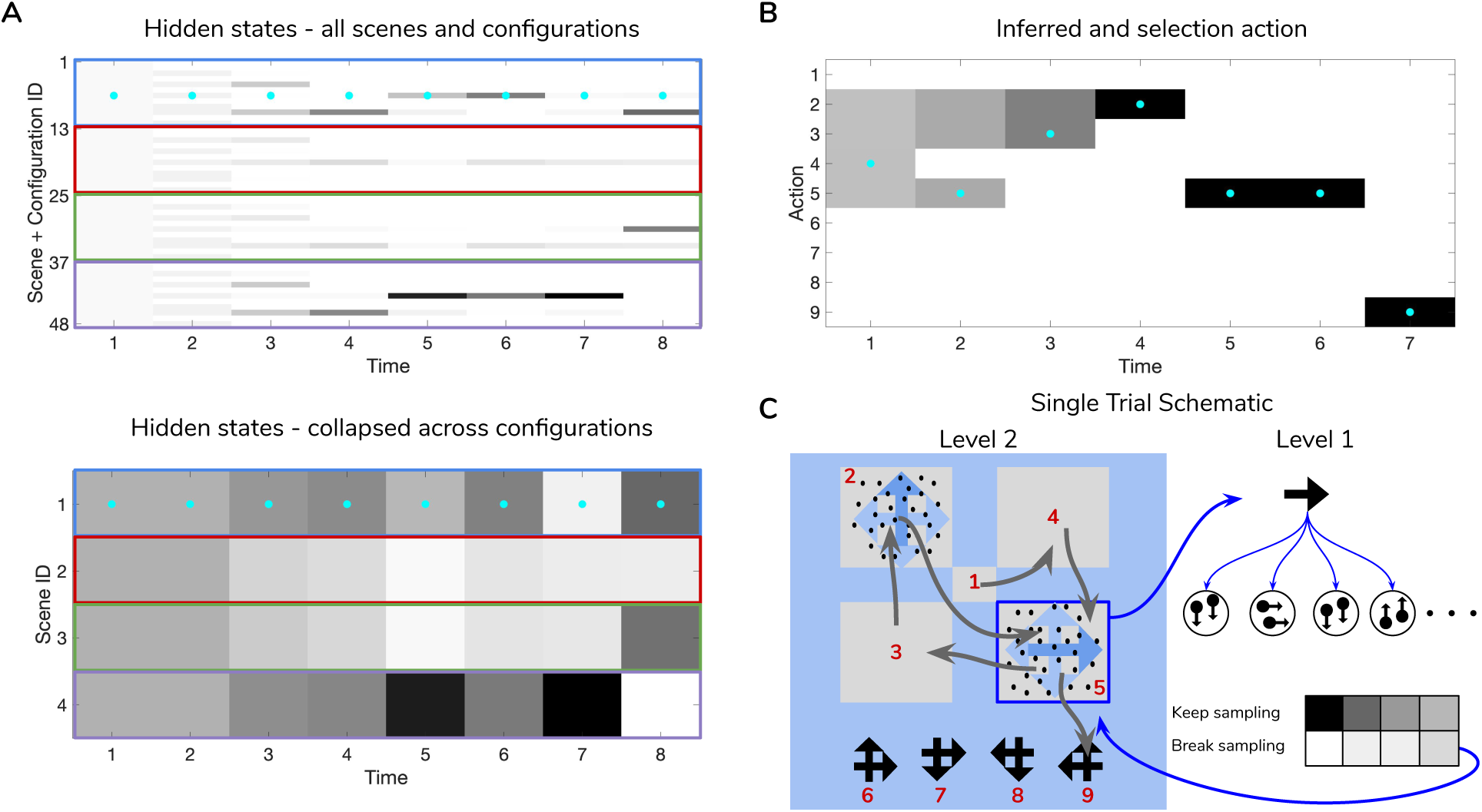
Simulated trial of scene construction with low sensory precision. Same as in Figure 9, except in this trial the precision of the mapping between RDM motion directions and samples thereof is lower, *p* = 0.5. This leads to an incorrect sequence of inferences, where the agent ends up believing that the scene identity is **LEFT-UP** and guessing incorrectly. Note that after this choice is made and incorrect feedback is given, the agent updates their posterior in terms of the ‘next best’ guess, which is from the agent’s perspective either **UP-RIGHT** or **DOWN-LEFT** (see the posterior at Time step 8 of Panel **A**). Panel **C** shows that the relative imprecision of the Level 1 likelihood results in a sequence of stochastic motion observations that frequently diverge from the true motion direction (in this case, the true motion direction is **RIGHT** in the lower right quadrant (Location 5). Level 1 belief-updating gives rise to an imprecise posterior belief over motion directions that are passed up as inferred outcomes to Level 2, leading to false beliefs about the scene identity. Note the ‘ambivalent,’ quadrant-revisiting behaviour, wherein the agent repeatedly visits the lower-right quadrant to resolve uncertainty about the RDM stimulus at that quadrant.

Figure 9 is a simulated trial of scene construction with sensory uncertainty at the lower level set to *p* = 5.0. This manifests as a stream of motion observations at the lower level that reflect the true motion state *∼* 98% of the time - i.e. highly-coherent motion. As the agent visually interrogates the 2 x 2 visual array (the 2^nd^ to 5^th^ rows of Panel **B**), posterior beliefs about the hidden scene identity (Panel **A**) converge on the true hidden scene. After the first RDM in the lower right quadrant is seen (and its state resolved with high certainty), the agent’s Level 2 posterior starts to only assign nonzero probability to scenes that include the **RIGHT**wards-moving motion stimulus. Once the second, **UP**wards-moving RDM stimulus is perceived in the upper left, the posterior converges upon the correct scene (in this case, indexed as state 7, one of the 12 configurations of **UP-RIGHT**). Once uncertainty about the hidden scene is resolved, **G** becomes dominated by instrumental value, or the dot-product of counterfactual observations with prior preferences. Expecting to receive correct feedback, the agent saccades to location 6 (which corresponds to the scene identity **UP-RIGHT**) and receives a ‘Correct’ outcome in the second-modality of Level 2 observations. The agent thus categorises the scene and remains there for the remainder of the trial to exploit the expected instrumental value of receiving ‘Correct’ feedback (for the discussion about how behaviour changes with respect to prior belief and sensory precision manipulations, we only consider behaviour up until the time step of the first categorization decision).

Figure 10 shows a trial when the RDMs are incoherent (*p* = 0.5, meaning the Level 1 likelihood yields motion observations that reflect the true motion state *∼* 35% of the time). In this case, the agent fails to categorise the scene correctly due to the inability to form accurate beliefs about the identity of RDMs at Level 1 - this uncertainty carries forward to lead posterior beliefs at Level 2 astray. Interestingly, the agent still forms relatively confident posterior beliefs about the scene (see the posterior at Timestep 7 of Figure 11**A**), but they are inaccurate since they are based on inaccurate posterior beliefs inherited from Level 1. This is because even though the low-level posteriors are built from noisy observations, the nonlinear, path-dependent nature of variational belief-updating means that the posterior ends up ‘focusing’ on a particular hidden state, which then gets integrated with empirical priors and subsequent observations to narrow the belief-space of possible hidden scenes. The hidden state that ends up receiving the most relative probability in the posterior depends on the stream of motion observations accumulated at the lower level, which are probabilistic samples from the corresponding column of **A**^(1),1^. The posterior uncertainty also manifests as the time spent foraging in quadrants before making categorization (nearly double the time spent by the agent in Figure 9). The cause of this increase in foraging time is two-fold. First of all, since uncertainty about the scene identity is high, the epistemic value of policies that entail fixations to RDM-containing quadrants remains elevated, even after all the quadrants have been visited. This is because uncertainty about hidden states is unlikely to be resolved after a single saccade to a quadrant with an incoherent RDM, meaning that the epistemic value of repeated visits to such quadrants decreases slowly with repeated foraging. Secondly, since Level 2 posterior beliefs about the scene identity are uncertain and are distributed among different states, the instrumental value of categorization actions remains low - remember that instrumental value depends not only on the instrumental value of receiving ‘Correct’ feedback, but also on the agent’s expectation about the *probability* of receiving this feedback upon making an action, relative to the probability of receiving ‘Incorrect’ feedback. The relative values of the prior preferences for being ‘Correct’ vs. ‘Incorrect’ thus tune the risk-averseness of the agent, and manifest as a dynamic balance between epistemic and instrumental value. See (Mirza, Adams, Parr, & Friston, 2019) for a quantitative exploration of these prior preferences and their effect on active inference.

**Figure 11:**
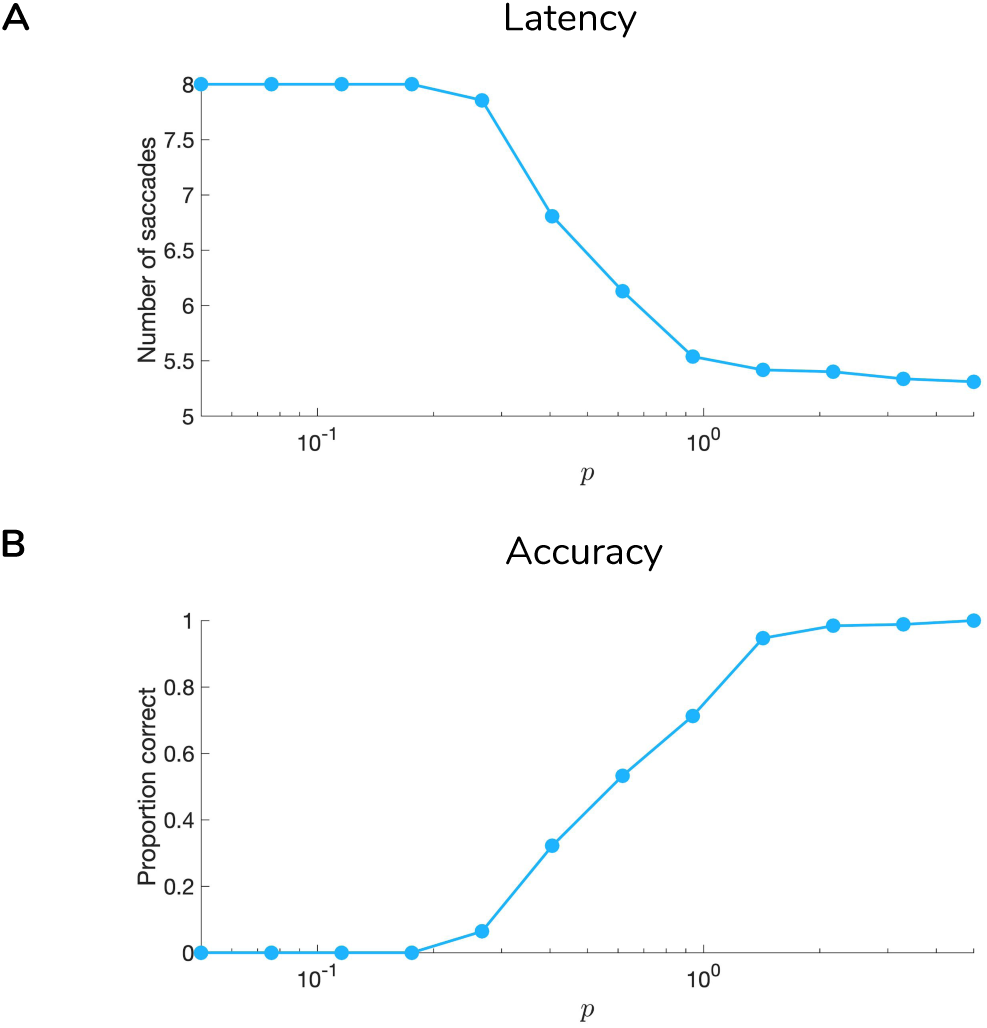
Effect of sensory precision on scene construction performance. Average categorization latency (**A**) and accuracy (**B**) as a function of sensory precision *p* which controls the entropy of the (Level 1) likelihood mapping from motion direction to motion observation. We simulated 185 trials of scene construction under hierarchical active inference for each level of *p* (12 levels total), with scene identities and configurations randomly initialised for each trial. Sensory precision is shown on a logarithmic scale.

We quantified the relationship between sensory precision and scene construction performance by simulating scene construction trials under different sensory precisions *p* (see Figure 11). The two measures shown are: 1) *categorization latency* (Figure 11**A**), defined as the number of time steps elapsed before a saccade to one of the choice locations is initiated; and 2) *categorization accuracy* (Figure 11**B**), defined as percentage of trials when the agent’s first categorization resulted in ‘Correct’ feedback. In agreement with intuition, for low values of *p* agents take more time to categorise the scene and categorise less accurately. As sensory precision increases, agents require monotonically less time to forage the array before categorizing, and this categorization also becomes more accurate. In the next section, we explore the relationship between sensory precision and performance when the agent entertains prior beliefs of varying strength about the probability of a certain scene.

### 7.2 Manipulating Prior Beliefs

For the simulations discussed in the previous section, agents always start scene construction trials with ‘flat’ prior beliefs about the scene identity. This means that the first factor of the prior beliefs about hidden states at Level 2 **D**^(2),1^ was initialised as a uniform distribution. We can manipulate the agent’s initial expectations about the scenes and their spatial arrangements by arbitrarily sculpting **D**^(2),1^ to have high or low probabilities over any state or set of states. Although many manipulations of the Level 2 prior over hidden states are possible, here we introduce a simple prior belief manipulation by uniformly elevating the prior probability of all spatial configurations (12 total) of a single type of scene. For example, to furnish an agent with the belief that there’s a 50% chance of any given trial being a **RIGHT-DOWN** scene, we simply boost the probabilities associated with hidden states 13 - 25 (the 12 spatial configurations of the **RIGHT-DOWN** scene) relative to the other hidden scenes, so that the total integrated probability of hidden states 13 - 25 is 0.5. This implies that the other hidden scenes each now have 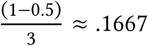 probability, once respectively integrated over their 12 configuration states. Figure 12 shows the effect of parametrically varying the strengths of prior beliefs on the same behavioural measures shown in Figure 11. Similar to Figure 11, Figure 12 demonstrates a monotonic increase in accuracy with increasing sensory precision, regardless of how much the agent initially expects a particular scene type. This means that strong but incorrect prior beliefs (over initial states) can still be ‘overcome’ with reliable enough sensory data. However, agents with stronger priors are less sensitive to the increase in sensory precision than their ‘flat-priored’ counterparts, as can be seen by the lower accuracy level of the most purple-colored lines in Figure 12. Note that the averages shown are only for agents with ‘incorrect’ prior beliefs; namely, the prior over hidden states in the generative model for each trial was always initialised to be a different scene type than the true scene. This has the effect of setting the minimum accuracy for the ‘strongest-priored’ agents (who typically categorise the scene identity at the first time step) at 0% rather than 25% (chance performance). These results are consistent with the fundamental relationship between the likelihood term and prior probability in Bayes’ theorem (see Equation (3)): the posterior over hidden states is calculated as the product of the likelihood and the prior. Increasing the precision of one of these two will ‘shift’ the posterior distribution in the respective direction of the more precise distribution. This manifests as a parametric ‘de-sensitizing’ of posterior beliefs to sensory evidence as priors become stronger. This balance between sensory and prior precision is exactly manifested in the prior-dependent sensitivity of the accuracy curves in Figure 12**B**.

**Figure 12:**
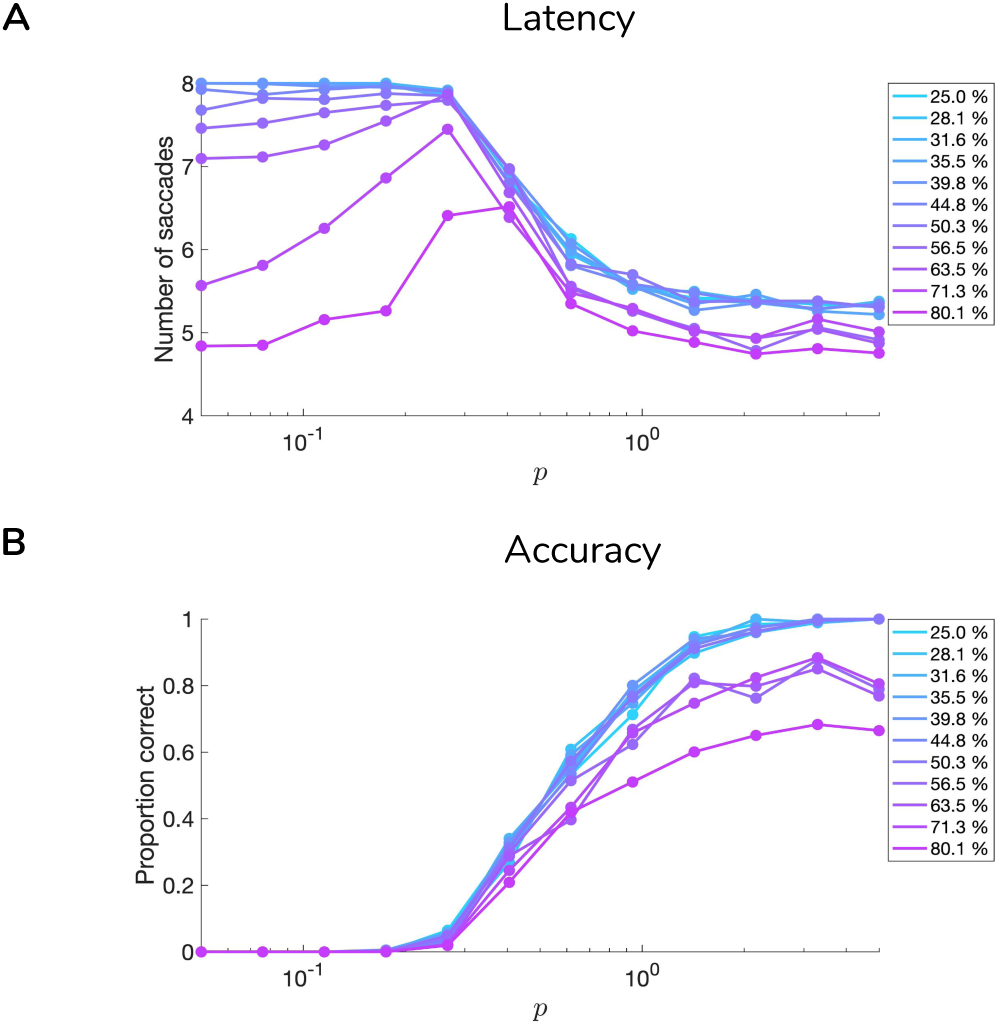
Effect of sensory precision on scene construction performance for different prior belief strengths. Same as in Figure 11 but for different strengths of initial prior beliefs (legend on right). Prior belief strengths are defined as the probability density of the prior beliefs about hidden states (1^st^ hidden state factor of Level 2 - **D**^(2),1)^) concentrated upon one of the four possible scenes. This elevated probability is uniformly spread among the 12 hidden states corresponding to the different quadrant-configurations of that scene, such that the agent has no prior expectation about a particular arrangement of the scene, but rather about that scene type in general. Here, we only show the results for agents with ‘incorrect’ prior beliefs - namely, when the scene that the agent believes to be at play is different from the scene actually characterizing the trial.

The interaction between sensory and prior precision is not as straightforward when it comes to categorization latency. Figure 12**A** shows that when the sensory precision *p* is high enough, most of the variance in latency introduced by prior beliefs vanishes, since observations alone can be relied on to ensure fast inference about the scene. For low values of *p*, however, latency is highly-sensitive to prior belief strength. Under weak prior beliefs and low *p*, the agent displays ambivalence - beliefs about RDM direction at Level 1 are not precise enough to enable scene inference, causing the agent to choose the policies that have (albeit) small epistemic value while avoiding the risk of categorizing incorrectly. This causes the agent to saccade among RDM-containing quadrants. Agents with stronger prior beliefs, however, do not rely on observations to determine posterior beliefs because their prior beliefs about the scene already lend high instrumental value to categorization actions. This corresponds to trials when the agent categorises the scene immediately (for the strongest prior beliefs, this occurs even before inspecting any quadrants) and relies minimally on sensory evidence. This faster latency comes at the cost of accuracy, however, as evident from the lower average accuracy of strongly-priored agents displayed in Figure 12**B**.

Now we explore the effects of sensory and prior precision on belief-updating and policy selection at the lower level, during a single quadrant fixation. Figure 13**A** shows the effect of increasing *p* on the break-time (or to analogise it more directly to eye movements: the fixational ‘dwell time’) at Level 1. We observe a nontrivial, inverted-U relationship between the logarithm of *p* (our analog of motion coherence) and the time it takes for agents to break the sampling at Level 1. For the lowest (most incoherent) values of the likelihood precision *p*, the agents dwell for as little time as they do as for the highest precisions. Understanding this paradoxical effect requires a more nuanced understanding of epistemic value. In general, increasing the precision of the likelihood mapping increases the amount of uncertainty that observations can resolve about hidden states, thus lending high epistemic value to policies that disclose such observations (Parr & Friston, 2017b). An elevated epistemic value predicts an increase in dwell time (i.e. via an increase in the epistemic value for the ‘**Keep-sampling**’ policy at Level 1) for increasing sensory precision. However, an increased precision of the Level 1 likelihood *also* implies that posterior uncertainty is resolved at a faster rate (due to high mutual information between observations and hidden states), which suppresses epistemic value over time. The rate at which epistemic value drops off thus increases in the presence of informative observations, since the posterior converges to a tight probability distribution relatively quickly. On the other hand, at very low likelihood precisions, the low information content of observations in addition to the linearly-increasing cost of sampling (encoded in the Level 1 preferences **C**^(1),2^) renders the sampling of motion observations relatively useless for agents, and it ‘pays’ to just break sampling early. This results in the pattern of break-times that we observe in Figure 13**A**.

**Figure 13:**
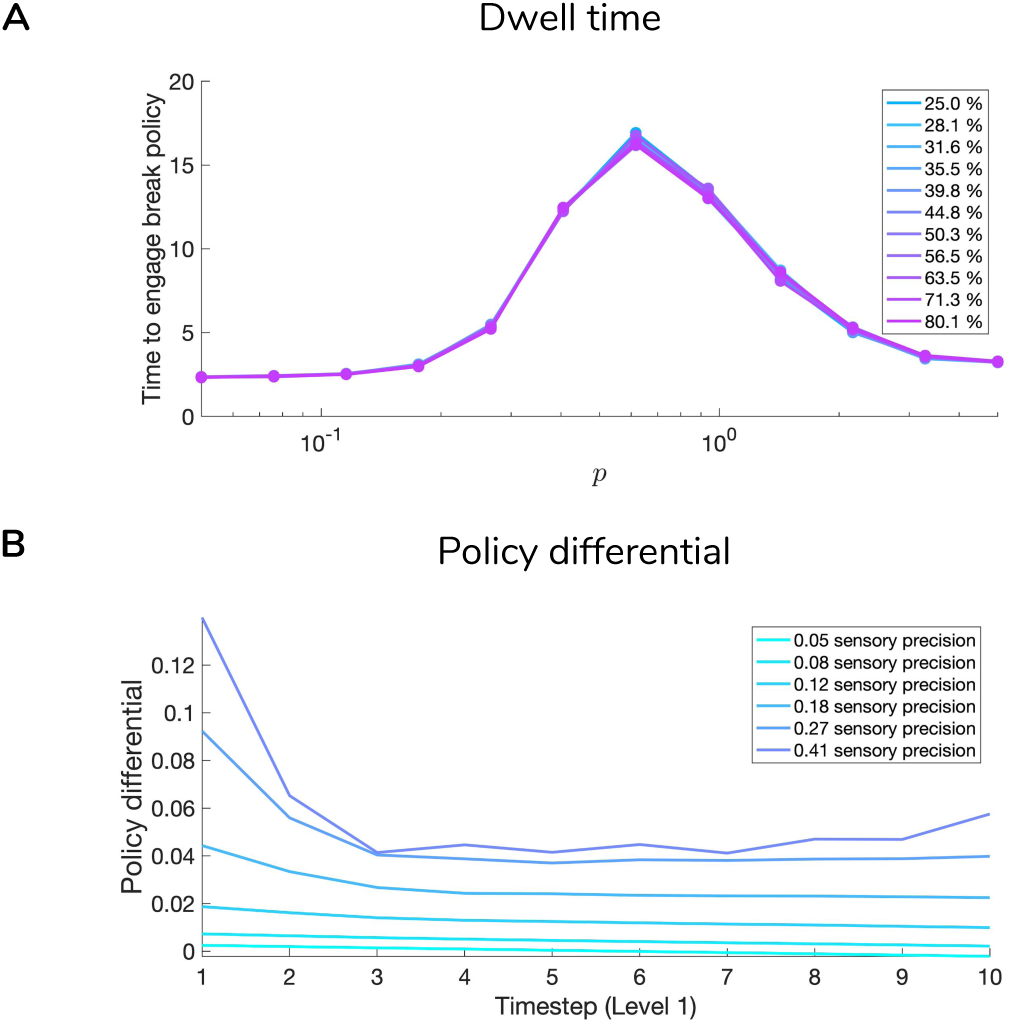
Effect of sensory precision on quadrant dwell time. Panel **A** shows the effect of increasing sensory precision at Level 1 on the time it takes to switch to ‘**Break-sampling**’ policy. Here, 250 trials were simulated for each combination of sensory precision and prior belief strength, with priors over hidden states at Level 2 randomly initialised to have high probability about 1 of the 4 scene types. Break-times were analyzed only for the first saccade (at Level 2) of each trial. Panel **B** shows the effect of sensory precision on evolution of the relative posterior probabilities of the ‘**Keep-sampling**’ vs. the ‘**Break-sampling**’ policies (*Policy Differential = P*_*Keep-sampling*_ *− P*_*Break-sampling*_). We only show these posterior policy differentials for the first 10 time steps of sampling at Level 1 due to insufficient numbers of saccades that lasted more than 10 time steps at the highest/lowest sensory precisions (see Panel **A**). Averages are calculated across different prior belief strengths, based on the lack of an effect, as is apparent in Panel **A**. The policy differential defined in this way is always positive because as soon as the probability of ‘**Break-sampling**’ exceeds that of ‘**Keep-sampling**’ (i.e., *Policy Differential* < 0), the ‘**Break-sampling**’ policy will be engaged with near certainty. This is due the high precision over policies at the lower level (here, *γ* = 512), which essentially ensures that the policy with higher probability will always be selected.

It is worth mentioning the barely noticeable effect of prior beliefs (Figure 13**A**) about the scene identity on break times at Level 1. Although prior beliefs about the scene at Level 1 manifest as empirical priors over hidden states (motion directions) at Level 2, it seems that the likelihood matrix plays a much larger role in determining break times than the initial beliefs. This means that even when the agent initially assigns relatively more probability to particular RDM directions (conditional on beliefs about scenes at Level 2), this initial belief can quickly be revised in light of incoming evidence (namely, observations at Level 1, inverted through the likelihood mapping to produce a marginal posterior over hidden states). This also speaks to the segregation of belief-updating between hierarchical levels; although beliefs about hidden states and observations are passed up and down the hierarchy, belief-updating occurs only with respect to the variational free energy of a particular layer’s generative model, thus insulating variational updating to operate at distinct spatiotemporal scales. This results in the conditional independence of decision-making across hierarchical levels, and clarifies the dissociable influence of prior about scenes on Level 1 vs. Level 2. For example, even on trials when an agent has strong prior beliefs about the scene and thus takes fewer saccades to categorise it, differences in lower-level ‘dwell time’ are still largely determined by the sensory precision *p* of the likelihood mapping and the preference to enter the ‘**Break-sampling**’ state, encoded as an increasing probability to observe oneself occupying this state (in **C**^(1),2^).

The curves in Figure 13**B** clarify the rate at which epistemic value decreases for high sensory precisions. The ‘policy differential’ measures the difference between the posterior probability of the ‘**Keep-sampling**’ versus ‘**Break-sampling**’ policies at Level 1: *P*Keep-sampling− *P* Break-sampling. At the lowest sensory precisions, there is barely any epistemic value to pursuing the ‘**Keep-sampling**’ policy, allowing the break policy to increasingly dominate action-selection over time. For higher sensory precisions, the ‘**Keep-sampling**’ policy starts with > 10% more probability than the ‘**Break-sampling**’ policy since the epistemic value of sampling observations is high, but quickly loses its advantage as posterior uncertainty is resolved. At this point the probability of breaking becomes more probable, since posterior beliefs about the RDM are fairly resolved and the instrumental of breaking is only getting higher with time.

## 8 Discussion

In the current work, we presented a hierarchical Markov Decision Process model of scene construction, where scenes are defined as arbitrary constellations of random dot motion (RDM) stimuli. Inspired by an earlier model of scene construction (Mirza et al., 2016, 2018) and a deep temporal formulation of active inference (Friston, Rosch, et al., 2017), we cast this scene construction task as approximate Bayesian inference occurring across two hierarchical levels of inference. One level involves optimizing beliefs about the instantaneous contents of agent-initiated visual fixations; the second level involves integrating the contents of different fixated locations to form beliefs about a higher-level concept like a scene. Through simulations we showed how this deep, temporal model formulation can be used to provide an active inference account of behaviour in such compositional inference tasks. Deep active inference agents performing scene construction exhibit the Bayesian hallmarks of a dynamic trade-off between sensory and prior precision when it comes to scene inference and saccade selection. The hierarchical segregation of inference between saccadic and fixational levels gives rise to unexpected effects of sensory uncertainty at the level of single fixations, where we observe an inverted-U relationship between motion coherence and fixational dwell time. This nonlinear relation can be explained by appealing to the evolution of epistemic value over time, under the assumption that the agent entertains beliefs about the precision of the environmental process generating visual sensations, while simultaneously optimizing the sufficient statistics of beliefs about the currently-fixated stimulus. The fact that the precision of the likelihood mapping increases the epistemic value of policies that furnish observations sampled from the generative process, while simultaneously increasing the rate at which posterior uncertainty is reduced, explains the non-monotonic influence of sensory precision on Level 1 decision latency.

These results contrast with the predictions of classic evidence accumulation models like the drift-diffusion model or DDM (Ratcliff, 1978; Palmer et al., 2005; Ratcliff & McKoon, 2008b). In the drift-diffusion model, reaction times are modeled as proportional to the latency it takes for a time-varying decision variable (or **DV**) to reach one of two fixed decision boundaries **Z** and −**Z** that respectively correspond to two hypotheses (e.g., the equivalent of sufficiently-strong posterior beliefs in one of two hidden states). At each time step, increments to the **DV** are calculated as the log of the ratio between the evidence for each hypothesis conditioned on observations. In discrete-time environments this update-rule for **DV** is equivalent to the Sequential Probability Ratio Test formulated by (Wald, Wolfowitz, et al., 1948). For time-independent decision boundaries and a fixed initial value of the **DV**, a drift-diffusion process yields a monotonic decreasing relationship between motion incoherence and decision latency (Bogacz et al., 2006; Ratcliff & McKoon, 2008a), where motion coherence factors into the DDM as the drift rate of the **DV** - this is analogous to the *sensitivity* of the **DV** to incoming sensory evidence. In the current active inference model, we have binarised policies at Level 1 in part to invite comparison between our model and DDM models (which in their classical form handle binary hypotheses). Rather than modelling actions as discrete perceptual decisions about the most likely hidden state underlying observations (since in the current context, we have a 4-dimensional RDM state space), we instead model the decision as the selection one of two ‘sampling’ policies, whose probabilities change over time due to the dynamics of the expected free energy. This evolving action-probability weighs epistemic drives to resolve uncertainty against prior preferences that encode an increasing ‘urgency’ to break sampling. This parameterization of decision-making permits a flexible (and in this case, somewhat unexpected) relationship between sensory uncertainty and decision latency (see Figure 13). We thus provide a novel, principled prediction for the relationship between sensory uncertainty and reaction time at different levels of inference in perceptual decision-making tasks.

In future work, we plan to estimate the parameters of hierarchical active inference models using data collected from human participants performing a scene construction task in which the identities of visual stimuli are uncertain (the equivalent of manipulating the sensory likelihood at Level 1 of the hierarchy). We propose our hierarchical active inference model to be a generic free-energy formulation of active sensing and evidence accumulation occurring in hierarchically-structured but inherently noisy environments.

## Acknowledgements

The authors would like to thank Brennan Klein for extensive feedback on the paper and the figures, and Brennan Klein and Alec Tschantz for discussion and initial conceptualization of the project. The authors also thank Kai Ueltzhöffer for feedback on the project and discussions relating active inference to drift-diffusion models. The authors also thank the Monash University Network of Excellence for supporting the workshop ‘Causation and Complexity in the Conscious Brain’ (Aegina, Greece 2018) at which many of the ideas related to this project were developed.

## Author contributions statement

RCH and MBM conceived the hierarchical active inference model. RCH, AP, and IK designed the scene construction task based on random dot motion stimuli. MBM, TP and KF gave critical insight into formulation of the model. RCH conducted the simulations and analyzed the results. All authors contributed to the writing of the manuscript.

## Data availability and software

The data used in this study are the results of numerical simulations, and as such, we do not provide datasets. The software used to simulate the data and generate associated figures can be obtained by the authors upon request, and will be included in an upcoming release of SPM12, which can be freely downloaded from https://www.fil.ion.ucl.ac.uk/spm/ as part of the DEM toolbox).

## Footnotes

1 From now on we assume the use of discrete probability distributions for convenience and compatibility with the sort of generative models relevant to the current work.

2 The Kullback-Leibler divergence or *relative entropy* is a non-negative measure of dissimilarity between probability distributions.

3 Hereafter we refer to observations and *hidden states* as *o* and *s*, respectively. We use the more generic term *hidden causes x* to refer to *all* aspects of the posterior - including hidden states, policies, and hyperparameters of the generative model.

4 This threshold is referred to as ‘residual uncertainty’, and by default is set to as 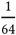 nats.

